# Machine Learning-Guided Engineering of High-Affinity Cross-Reactive Antibodies with Minimal Mutations

**DOI:** 10.64898/2026.07.15.738714

**Authors:** Hugo Dorison, Anne-Laure Grindel, François Thenier, Chloé Pluchart, Mélanie Munch, Càtia Oliveira, Steven Dubois, Camille Le Drezen, Raphaël Guérois, Bernard Maillère, Charles Truillet, Hervé Nozach

## Abstract

Antibodies raised against human targets often fail to recognize their animal orthologs, limiting preclinical evaluation in relevant models. We developed a Deep Mutational Scanning (DMS)-coupled deep learning strategy to engineer potent cross-reactive antibodies with minimal sequence divergence. Starting from C4, a fully human anti-PD-L1 antibody with weak recognition of murine PD-L1, DMS identified substitutions that improved binding to both human and mouse antigens. Conventional recombination of beneficial mutations generated highly cross-reactive antibodies but required 13 to 15 substitutions. To reduce this mutational burden, a deep learning model trained on DMS-derived sequence-binding data was used to identify minimal mutation combinations predicted to retain high affinity. This approach yielded variants carrying only 4 to 5 substitutions, with *in vitro* and cellular binding properties comparable to highly mutated antibodies. Epitope mapping, structural modeling and *in vivo* assessment further confirmed that these engineered antibodies retained PD-1/PD-L1 blockade and demonstrated therapeutic activity in a mouse tumor model.

## Main text

Antibodies are widely valued for their ability to bind to single targets with high affinity and selectivity, while being relatively easy to produce [1]. However, for therapeutic antibodies, the transition from preclinical to clinical development remains a major bottleneck, in part because species-specific differences can limit the predictive value of preclinical efficacy studies. Indeed, antibodies directed against human proteins often display limited cross-reactivity with their animal counterparts, owing to substantial evolutionary divergence between the orthologous targets. Such stringent target specificity can impede meaningful *in vivo* evaluation of biological activity and therapeutic potency, and may therefore represent a significant limitation during drug development. Consequently, *in vivo* assessments are sometimes carried out using surrogate antibodies [2,3] but these studies provide only limited insight into the *in vivo* biological effects of the initial therapeutic antibody. In fact, even minor differences in affinity between human and animal targets may hinder the translational relevance of animal model studies. Alternatively, genetically engineered animal models can be generated to test a specific therapeutic antibody [4,5] but expression of the transgene may not fully recapitulate the natural endogenous expression [6,7]. Together, these limitations highlight the value of cross-species antibodies that recognize both the human target and its relevant animal ortholog.

Cross-reactive antibodies can be generated by using antibody display systems, such as phage and yeast display, to select variants that recognize orthologous targets across species. These approaches rely on a physical linkage between the antibody-encoding sequence and the displayed antibody, together with the generation of mutational libraries that are screened for antigen recognition. The binders are then sorted and identified by DNA sequencing. To increase the likelihood of identifying hit candidates, mutational libraries were initially designed to maximize diversity. Successful campaigns often yield highly mutated candidates [8,9], with mutational distance from the parental sequence often viewed as a marker of higher potential affinity and thus as a positive trait [10]. However, a high mutational load is not necessarily optimal. While these substitutions often increase affinity, they may also impair other properties such as expression or stability [11]. Indeed, divergence from human germline sequences is often considered undesirable, prompting multiple groups to develop germinalization strategies aimed at optimizing antibody function while keeping sequences as close as possible to the human germline repertoire [12,13].

Indeed, the affinity-maturation process is unlikely to preserve germline residues and therefore generally decreases the sequence identity to human germline sequences, also referred to as antibody humanness [14]. This decrease in humanness may increase the risk of unwanted immune responses, further reinforcing the need to limit unnecessary sequence divergence. Although some substitutions may play a key role in affinity enhancement, not all selected mutations necessarily contribute directly to antigen binding. Some may instead improve properties specific to the *in vitro* selection system, such as display or solubility, whereas others may behave as passenger mutations carried along with affinity-enhancing substitutions.

In contrast, several mutagenesis campaigns have achieved significant affinity gains with only a few mutations [8–10,15], suggesting that many mutations can be dispensable. Identifying the minimal set of beneficial mutations required to improve binding is likely to depend on the initial affinity and the targeted epitope, while deciphering the contribution of substitutions embedded within multiple others remains challenging, even with modern next-generation sequencing, due to the complexity of mutational couplings.

In this study, we propose to leverage Machine Learning (ML), concomitantly with a Deep Mutational Scanning (DMS) process, to learn the sequence determinants underlying low, medium or high affinity for a selected antigen. New sequences can therefore be generated *in silico* and their effects predicted by the model. We applied our strategy to confer cross-reactivity to a fully human anti-PD-L1 antibody called C4 [16], specific to human PD-L1 that originally showed very limited binding to the murine ortholog. PD-L1 is overexpressed in malignant cancer cells to escape T cell targeting through PD-1/PD-L1 interaction [17,18]. Blockade of PD-1 binding to PD-L1, restores T cell negating the proliferating effect of PD-L1 overexpression in tumors and yielding antitumor immunity [19,20]. Several trademarked antibodies (Avelumab, Atezolizumab and Durvalumab) recognize PD-L1, confirming its relevance in oncology. All these antibodies recognize very similar overlapping epitopes, yet Durvalumab does not recognize the murine antigen [21–25], highlighting the challenge of obtaining cross-reactive antibodies. Our objective is to engineer C4 with sub-nanomolar affinity and cross-species reactivity toward both human and mouse PD-L1, in order to enable its preclinical assessment in relevant immunocompetent mouse models.

Since the parental C4 antibody displayed nanomolar affinity for the human antigen together with a weaker but measurable affinity for the murine ortholog, in the 200-300 nM range, we pursued an affinity maturation strategy designed to rebalance its affinity profile for human PD-L1 and its murine counterpart. As a first step, we performed conventional affinity maturation by combining DMS-identified beneficial mutations to generate a first-generation panel of molecules with robust cross-reactivity. However, because variants generated through the DMS process harbored 13 to 15 mutations, we next trained our ML model on the DMS dataset to identify combinations of substitutions that could preserve cross-reactivity while reducing mutational load. This ML-guided redesign enabled the engineering of novel variants carrying only 4 to 5 substitutions, with *in vitro* and *in vivo* properties indistinguishable from those of the highly mutated first-generation molecules. We further used state-of-the-art deep learning-based structural modeling to interpret the potential effects of four selected mutations. More broadly, this approach could be applied to other antibody engineering objectives requiring fine control of affinity, selectivity, or cross-reactivity while limiting mutational burden.

## Results

### Generation of high affinity cross-reactive antibodies against PD-L1

To precisely identify key paratope residues involved in human PD-L1 antigen recognition and determine which of them could be substituted to accommodate the murine antigen, a deep mutational scanning (DMS) approach was employed. Fab fragments corresponding to the C4 antibody were expressed on the surface of yeast cells and DMS was performed as described in [26] with adjustments. We generated a single-mutant library across the variable parts of the light and heavy chains of the C4 parental antibody with SOE-PCR and NNK codons (Fig 1A left). All substitutions were considered at each position, resulting in a manageable diversity of ≈ 4.10^3^ codons for the variable heavy chain (VH). The library of yeast cells displaying antibody fragments was sorted by Fluorescence-Activated Cell Sorting (FACS) to isolate clones exhibiting enhanced recognition of either the murine or human antigen compared to the parental antibody (yellow population, Fig. 1A, middle). Both the sorted and the corresponding unsorted libraries were sequenced to provide the basis for enrichment analysis. In order to achieve cross-reactivity, affinity for mouse PD-L1 must be increased, while preserving binding to human PD-L1. For that purpose, multi-layer heatmaps were generated by integrating all conditions to identify the hotspots associated with increase or decrease binding, depending on whether the libraries were selected for improved or impaired binding, respectively (Fig. 1A right).

**Figure 1:**
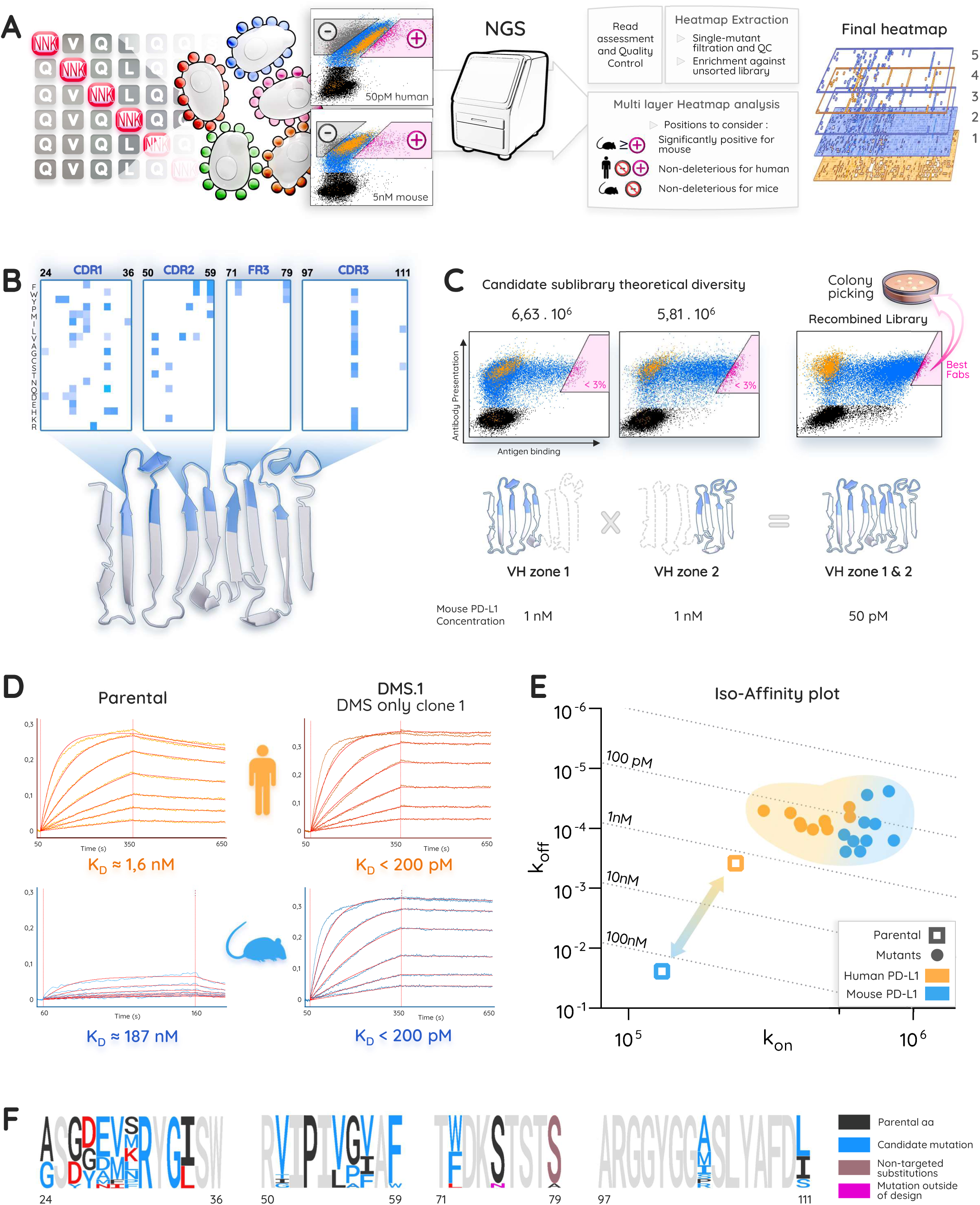
Deep Mutational Scanning (DMS) based affinity maturation to confer cross-reactivity to C4 antibody. (A) Schematic representation of the DMS process. (Left) A library of single-mutants was transformed into yeast cells through gap repair. (Middle) This library containing every substitution at each position of the VH was incubated with either human or mouse PD-L1 with a single biotin fusion (on the X axis of the cytograms: antigen binding, measured by the fluorescence of Strepatividin-PE bound to PD-L1. On the Y axis: antibody presentation, measured by APC fluorescence from an anti-V5 tag antibody recognizing a V5 tag on the Fab VL). A clonal parental sequence carrying a GFP reporter was spiked in the single-mutant population (orange population). Two gates were drawn per conditions to recover single mutants with a positive (pink) or negative (grey) effect on affinity in comparison with the parental clone. (Right) NGS readout and data analysis of the 4 populations recovered leading to multi-layers heatmaps: 1, unaltered heatmap on human; 2, unaltered heatmap on mouse; 3, significantly positive substitutions on mouse; 4, enriched positions on human; 5, resulting heatmap with compatible positions. (B) Multi-layer heatmap resulting from the analysis of VH variants. Candidate positions were selected using a structural model of the parental Fab with a focus on residues potentially oriented toward the antigen. (C) Substitutions identified in panel B were recombined in two lower-diversity sub-libraries and sorted independently. Selected populations from VH zones 1 and 2 were then recombined into a full VH library subjected to a second round of selection, and plated for random colony picking. (D) Representative BLI graphs used in panel E to plot each point. Orange curves: human PD-L1 measurements (two top graphs). Blue curves: mouse PD-L1 (two bottom graphs). (E) Iso-affinity plotting of 10 candidates randomly picked from the final candidate library. BLI measurements (K_D_) are plotted as the ratio of k_off_ and k_on_. Higher and to the right is better. Dotted lines: iso-affinity values. Orange and blue indicate biding to human and mouse PD-L1, respectively. The leftmost arrow indicates the large difference between the parental K_D_ values whereas all selected candidates cluster within a region of overall higher affinity. (F) Sequence logo of Sanger sequencing of 20 candidates obtained by random colony picking in the final candidate library.

To further enhance affinity, we next sought to combine favorable mutations within combinatorial libraries encompassing the VH domain. We selected mutations simultaneously exhibiting at least a 2-log enrichment for mouse PD-L1, while showing neutral or enhanced binding to human PD-L1 (Fig. 1B, Sup. Fig. 1). Candidate positions were then selected without a priori by using an AlphaFold3 model of C4 [27], in which all residues potentially oriented toward the antigen are highlighted in blue (Fig. 1B). These zones are centered on CDRH1, H2, H3 and FWK3 and span across 20 positions where selected substitutions are introduced while systematically including the parental amino acid. The VH combinatorial library was split into two sub-libraries to ensure each sub-library could be sampled at least 5 times within a single yeast transformation (Fig. 1C). The same workflow was applied to the C4 VL (Sup. Fig. 2), but the resulting combinatorial libraries showed minimal improvement in FACS and was not pursued further.

**Figure 2:**
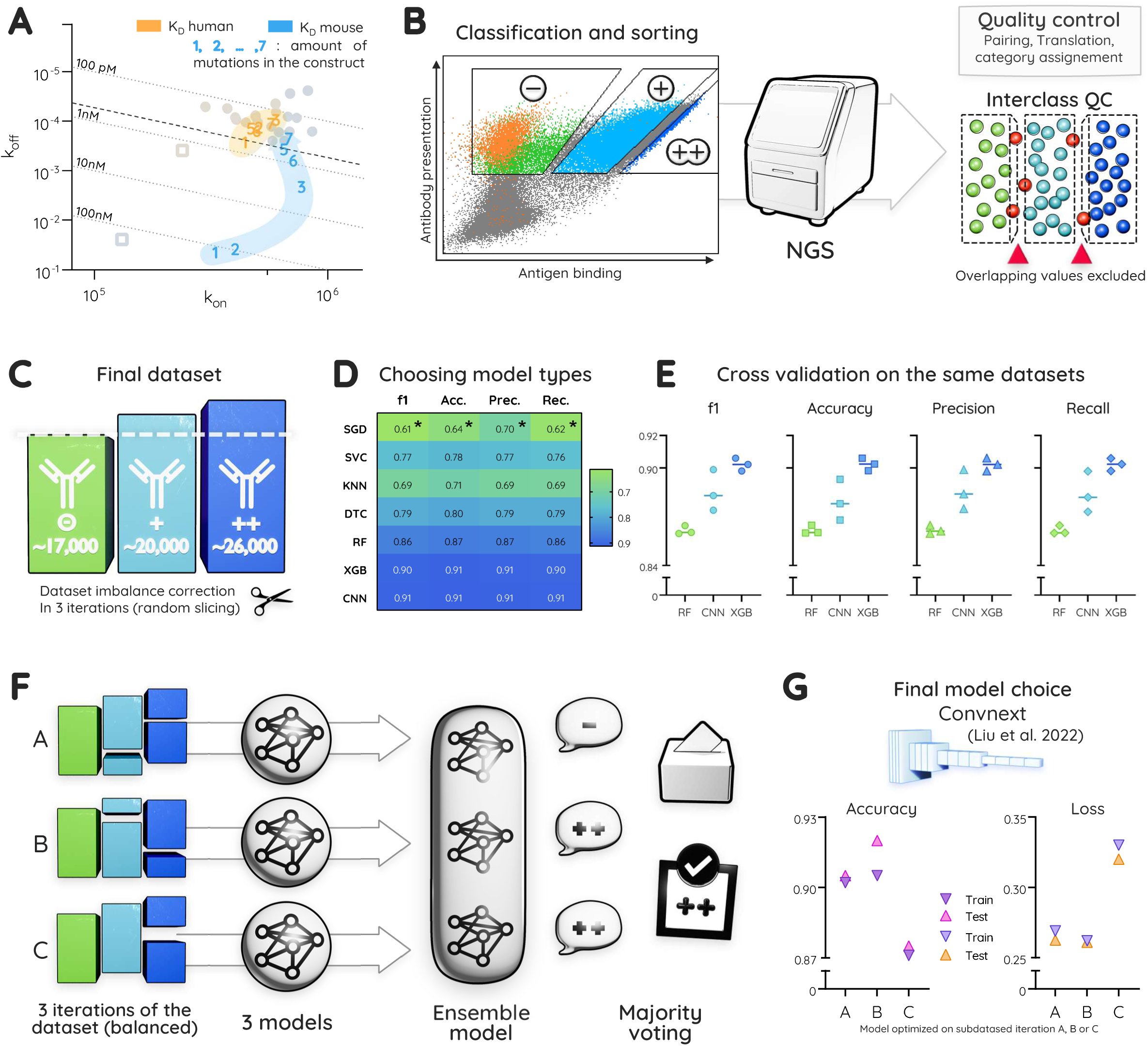
empirical and ML-guided approaches for generation of low mutations high affinity antibodies. (A) Iso-affinity plotting BLI measurements of IgGs generated by hand, through iterative additions of mutations starting from the parental C4. Overlayed on top of the iso-affinity plots from figure 1 (greyed out). Dotted line: lowest K_D_ value recorded from the DMS colony-picked clones. (B) ML workflow experimental design. (Left) Sorting strategy on final candidate sublibrary (rightmost panel fig 1.C) Orange: parental Fab displayed on GFP+ yeast. 3 gates drawn and sorted then independently read in NGS. (Right) Classic bioinformatic readout and Quality Control (QC) of the sequences followed by class assignment. Additional inter-class QC: sequences common to one or more classes are excluded from the analysis. (C) Final dataset after QC. Numbers indicates the amount of distinct sequences. Imbalance corrected to 17000 sequence per category through random exclusion of sequences from supernumerary classes. Process repeated 3 separate times to get 3 slightly different datasets to use in panel F. (D) Performance assessment of main model architectures on the full dataset: no imbalance correction. Plotted values are a mean of 4 iteration of the learn / test full cycle. All values are within confidence interval of 95% (IC95) < 0,01. Asterisks for values where IC95 < 0,05. SGD: Stochastic gradient descent; SVC: support vector classifier; KNN: k nearest neighbor; DTC: Decision tree learning; RF: Random Forest; XGB: XGBoost; CNN: convolutional neural network (ConvNeXt). Acc.: Accuracy, Prec.: Precision, Rec.: Recall. (E) Performance metrics of the best 3 models from panel D during cross-validation on the very same 3 balanced datasets with the same train and test subsets split with a 75 – 25 % ratio. (F) Machine learning strategy. Final dataset replicated in 3 iterations only different through the random dataset imbalance correction. Each iteration split for train / test. 1 model trained per dataset. Best possible models for each dataset recovered and used as an ensemble model for prediction. Consensus prediction is selected as the final prediction. (G) Selected model type. Accuracy and loss for train and test of the best 3 models (A, B, C) on the 3 iterations of the dataset. Best performance obtained on epoch 9, 10 and 5 for A, B and C respectively.

Improved candidates of each VH sub-libraries were sorted at low murine PD-L1 concentrations (Fig. 1C). Resulting diversity of both sub-libraries was recombined into a final library spanning the entire VH domain, and subjected to an additional round of sorting at even lower murine PD-L1 concentrations. Twenty individual clones were randomly selected by colony picking and Sanger sequencing. Ten clones were then recombinantly expressed as IgG molecules in mammalian cells and their affinity profiles were determined using bio-layer interferometry (BLI). Affinity improvements were considerable, nearing a 1000-fold improvement over the parental affinity for mouse PD-L1 (Fig. 1D). Interestingly, affinity for human PD-L1 was also increased. All clones displayed very high and similar affinities for both human and mouse PD-L1 as shown by their clustering in the iso-affinity graph (Fig. 1E). As opposed to the very large difference between the K_D_ of parental C4 (squares) for mouse (blue) and human (yellow) PD-L1. Sequence analysis of the clones resulting from this affinity maturation strategy revealed a high number of substitutions relative to the parental sequence, ranging from 13 to 15 per clone. A sequence logo representation highlights that some residues are not shared between clones, while others are highly conserved across all clones (Fig 1F, Sup. Fig. 3A, 3C). Although they may appear equally important in this incomplete snapshot, we hypothesize that their contribution to affinity may not be equivalent. In order to build high affinity cross-reactive antibodies that do not diverge that far from the parental sequence, we therefore aimed at deciphering their impact on binding.

**Figure 3:**
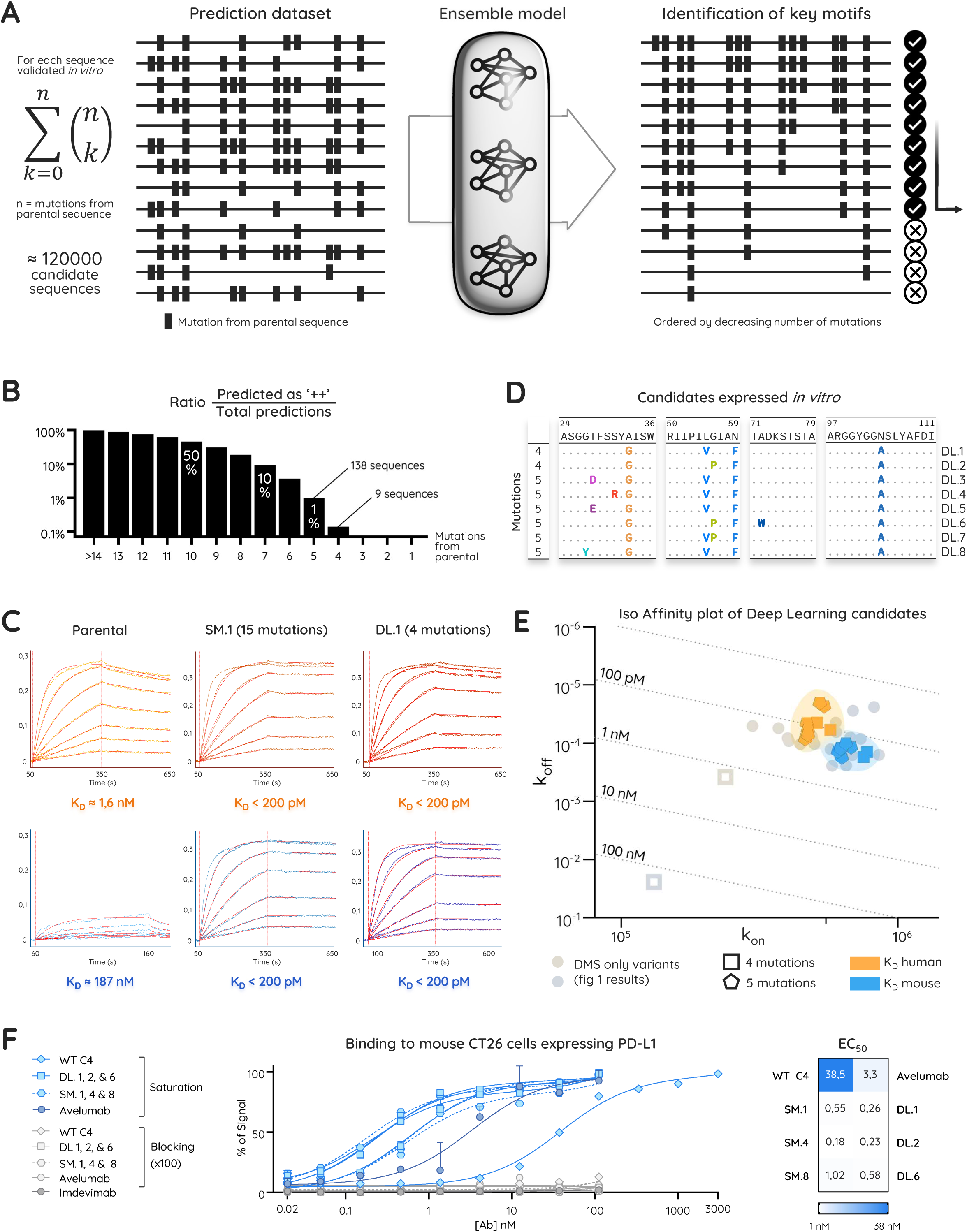
Prediction of minimal combinations of substitutions to reach high affinity cross-reactive antibodies. (A) Strategy for prediction of the least mutated sequence that will still perform *in vitro*. (Left) Generation of new combinations of mutations. Starting from the 10 sequences validated *in vitro* in fig. 1 panel D, substitutions from parental sequence are all removed one by one. Considering all possible permutations. Generation of new combinations of substitutions already encountered are generated. Typically n = 12 to 15. (Middle) sequences fed to ensemble model. Categories “-”, “+” and “++” are assigned to each sequence with a confidence metric. (Right) Prediction analysis: sequences predicted as “++” are filtered (marked with a checkmark (IZJ)) and sorted by decreasing number of mutations from parental. Target sequences are the least mutated in this filter (arrow). (B) Analysis of ensemble model predictions. All 120000 candidate sequences are split by number of mutations from parental. In Y axis is ploted the ratio of sequences predicted as “++” divided by the total number of sequences for a number of mutations. Since the starting sequences are the 10 validated *in vitro*, untouched sequences are more likely to be predicted as 100% “++”. No sequences are predicted as “++” under 4 mutations. (C) Excerpt of BLI data used to plot panel E. Top row: measurements on human PD-L1. Bottom row: mouse PD-L1. Column 1 and 2: parental IgG and candidate DMS.1 as in fig. 1 panel E. Column 3: DL.1 candidate generated from predictions. Vertical dotted line 1: start of association phase (160 or 300s): Anti-human capture tips covered in IgGs are dipped in antigen containing solutions. Dotted line 2: start of dissociation phase, tips are dipped in antigen-free solution. (D) Final candidates selected to synthetize and characterize *in vitro* as IgGs. 2 sequences from the 9 predicted as “++” with 4 substitutions from panel B. And 6 out of 138 possible sequences with 5 mutations. Top row: parental sequence with numbering on top. Candidates are displayed with only the substituted amino acids from parental. Consensus amino acids are shown with a dot. (E) Iso affinity plotting of BLI K_D_ affinity measurement of the 8 sequences in panel C. Greyed out in the bottom are the results from fig. 1 panel D for comparison. Higher and to the right is better. Dotted lines: iso-affinity values. Yellow and blue for affinity on human and mouse PD-L1 respectively. (F) Binding assay of the different IgG on CT26 mouse cells. Both saturation and blocking condition (by a 100 more concentrated unconjugated antibody) were tested in duplicates. Data are presented as mean ±SD. Data were fitted with a one-site binding model and were normalized for comparison, and EC50 were obtained (table on the right).

Our first effort focused on generating six new sequences carrying sparse, manually curated mutations based on expert analysis of Sanger sequencing data. Substitutions were introduced to the parental sequence one by one, starting with the ones we identified as most critical for affinity maturation (Sup. Fig. 3D). Surprisingly, the most widely conserved substitutions S31R, both alone and in combination with A33G, had no appreciable effect on K_D_ (Fig. 2A, Sup. fig. 3E). The inclusion of a third mutation, N59F, led to a substantial improvement in affinity, lowering the K_D_ to the single-digit nanomolar range. However, the lowest K_D_ recorded among our 10 colony-picked candidates was only reached by combining seven substitutions. While reducing the number of substitutions to seven is encouraging, this approach provided little insight into the effect of each mutation and it may be possible to achieve similar performance with even fewer changes. To address this, we devised a specific Machine Learning (ML)-based approach.

### Machine Learning (ML) can predict low, medium and high affinity binders

Instead of cherry-picking the very best candidates, we set out to explore the repertoire of candidates through deep sequencing. Library variants were clustered in 3 behaviors: “-“parental-like, “+” enhanced and “++” strongly enhanced antigen binding (Fig. 2B, left). The “++” gate is identical to the one used during our standard colony-picking strategy (Fig. 1C). The least mutated antibody identified within this gate harbored 8 substitutions, with an average of 13.8 mutations across the population. We thus chose to use a ML-based approach to design new, less mutated clones, *de novo*, by learning the properties of substitutions in different contexts. This setup enables the problem to be framed using discrete classes that facilitate ML, allowing the model to learn not only which substitutions are beneficial, but also which can be omitted across different sequence contexts. Sorted populations were deep sequenced. After standard quality control steps, single read sequences and the few sequences present in more than one gate were discarded (Fig. 2B, middle). A relatively well-balanced dataset was obtained with 17,000, 20,000 and 26,000 unique sequences in the “- “, “+ “and “++ “group, respectively (Fig. 2C).

Various ML architectures were evaluated over the whole dataset based on their ability to correctly classify unseen test sequences among the three gates, with higher f1 values indicating more accurate test predictions [28]. All tested architectures demonstrated performance indicative of genuine learning, with accurate predictions on the test datasets that were unseen during training. Among the architectures tested in this preliminary comparison (Fig. 2D), the three best-performing models were global models: a Random-Forest (RF) algorithm, XGBoost (XGB) and finally ConvNeXt [29], a Convolutional Neural Network (CNN). To ensure that performance was not dependent on a single arbitrary train/test partition, the three architectures were compared across three independent random train/test splits with fixed proportions (Fig. 2E). Results were consistent with preliminary assessments over the whole dataset for RF and XGB. The CNN reached high performance values in some runs but showed greater variability and slightly lower performance than XGB. Nevertheless, we selected the CNN for the downstream design step because it could be trained directly on the full antibody sequences, without restricting the input to the substitutions included in the library design. In contrast, the XGB model required a reduced representation focused on the designed mutation positions. The CNN therefore offered more flexible framework to capture serendipitous mutations outside of design, as observed prior in the sequence logos of the DMS clones (Fig. 1F). In addition, CNN performance can be further optimized with hyper-parameter tuning.

### Deep Learning can be used to generate new candidates with minimal mutations

We chose a straightforward one-hot encoding approach [30–32] for the amino acid sequences. To alleviate Dataset imbalance, we randomly removed sequences from the overrepresented groups to reach the smallest class size of 17,000 sequences. This was performed three times to make use of the dataset as much as possible. These datasets were used to train three independent models (Fig. 2C). Majority voting over the models was implemented for an ensemble-model strategy [30] to maximize the correlation between in silico predictions and *in vitro* results. The models were trained after empirical hyperparameter optimizations to reach satisfying accuracies of 0.875, 0.905 and 0.92 (Fig. 2G) on the test portion of the three datasets after about 10 epochs (Sup. Fig. 4).

**Figure 4:**
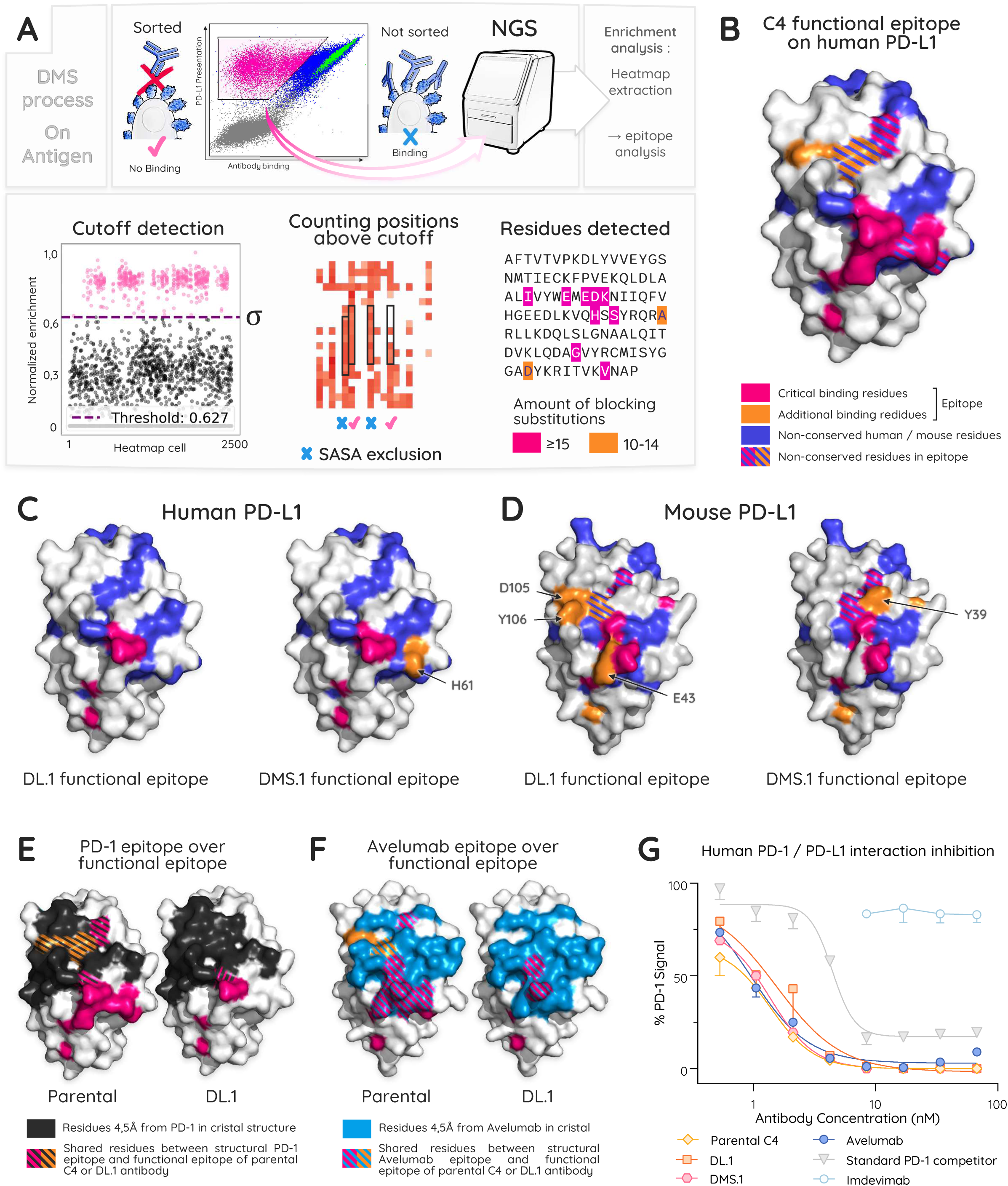
Deciphering functional epitopes and binding modes of parental and matured antibodies. (A) Experimental principle (Top) and analysis process (Bottom). (Top) DMS is reemployed as in fig. 1 panel A but single-mutations are inserted on the antigen sequence. PD-L1 (Nter domain) is now displayed at the surface of yeasts. Antibodies are the analyte revealed in the x axis of the cytogram through an alfa-tag fusion on the VL and a fluorescent anti-alfatag secondary antibody. As previously described, Y axis is HA detection through an anti-HA secondary antibody. The HA tag is fused to PD-L1 displayed on yeasts. Green: yeast displaying the wild-type PD-L1 non mutated. Cells of interest are the ones whose single-mutation inhibits the binding of PD-L1 to the target antibody: pink gate. The latter were sorted, sequenced (Novaseq) and enrichment analysis was performed against the unsorted library to obtain a Heatmap. (Bottom) Further analysis was needed to obtain the epitope: (Bottom left) All values of one heatmap are plotted to visualize the background noise from the residues highly engaged. Logs of enrichment are normalized per heatmap between 0 and 1 for comparison. Threshold is set at the standard deviation value of all the heatmap values: σ. (Bottom left) per residue on the parental sequence, the amount of substitutions above cutoff (between 0 and 20) are counted. Solvent accessible surface area (SASA) [59] is computed through pymol on an Alphafold 3 (AF3) prediction of the parental sequence and used to discard all residues for which SASA < 0,1. (Bottom right) Arbitrary coloration of residues for which over 15 substitutions abolish binding (pink): high importance in antibody binding. Residues for which 10 to 14 substitutions are deleterious are colored in orange: slightly less engaged residues. (B) Nter domain human PD-L1 from PDB entry 4ZQK [60] with parental antibody functional epitope. Pink and orange residues are engaged by the antibody on the antigen, they are the functional epitope. In blue: residues of PD-L1 that differ from human to mouse. Stripped blue pattern indicates residues that are in the epitope and different from human to mouse. (C) Functional epitopes of DL.1 and DMS.1 on human PD-L1. (D) Functional epitopes of DL.1 and DMS.1 on mouse PD-L1. Mouse PD-L1 Nter domain obtained through an Alphafold 3 prediction. (E) Overlay of the functional epitopes of Parental C4 (left) and DL.1 (right) with PD-1 structural epitope (black) on human PD-L1 (pdb: 3BIK [61]). (F) overlay of the functional epitopes of Parental C4 (left) and DL.1 (right) with Avelumab structural epitope (blue) on human PD-L1 [23]. (G) Competition ELISA between immobilized PD-L1, biotinylated PD-1 and antibodies to test. Imdevimab introduced as an irrelevant isotype control. Unknown reference PD-1 competitor antibody in grey provided by the ELISA kit manufacturer. Avelumab, a well characterized PD-L1 inhibitor [21] was introduced as a reference. Its epitope obtained by crystallography overlaps with functional epitopes of the parental C4 and engineered antibodies. As expected, it competes with PD-1 in this ELISA.

Our CNN ensemble model proved capable of assigning antibody affinity classes directly from sequence information, indicating that meaningful rules governing sequence-affinity relationships had been learned. To build the best possible compromise of high affinity and low mutation count, we sought to dissect these rules. We therefore aimed to identify the sequences with the fewest substitutions that would still be predicted as belonging to the highest-affinity class (“++”) by the ensemble model. To achieve this, we focused on determining which mutations were essential in the ten DMS-derived clones previously characterized (Fig. 1E). We generated an in silico dataset (Fig. 3A) by systematically removing substitutions one at a time and exploring all resulting permutations. Since the ten DMS derived sequences harbor more than 14 mutations, this resulted in a prediction dataset of 120,000 unique sequences, exceeding the size of the learning dataset. Each sequence was then classified into an affinity class by the ensemble model. Overall, removing mutations tended to shift the predicted classes to lower affinity ones. However, a subset of sparsely mutated combinations was still predicted to retain the highest-affinity class (Fig. 3B). According to our ensemble model, a minimum of four mutations was necessary to retain the highest affinity, and only nine sequences fit this criterion, representing 0.14% of the prediction dataset. We established a short-list of candidates to test *in vitro* (Fig. 3C, Sup. Fig. 5), selecting only sequences for which all models within the ensemble agreed, and for which confidence metrics were above 80%. Model decisions on sequences with 5 substitutions were unambiguous. Due to the substantially lower number of mutations, the candidates selected with four or five mutations exhibit humanness ratios of 95.9% and 94.9%, respectively, markedly higher than those of the highly mutated DMS-derived sequences (86–89%, Sup. Fig. 3B). We therefore experimentally validated the performance of these new sequences and compare them with the highly mutated variants.

**Figure 5:**
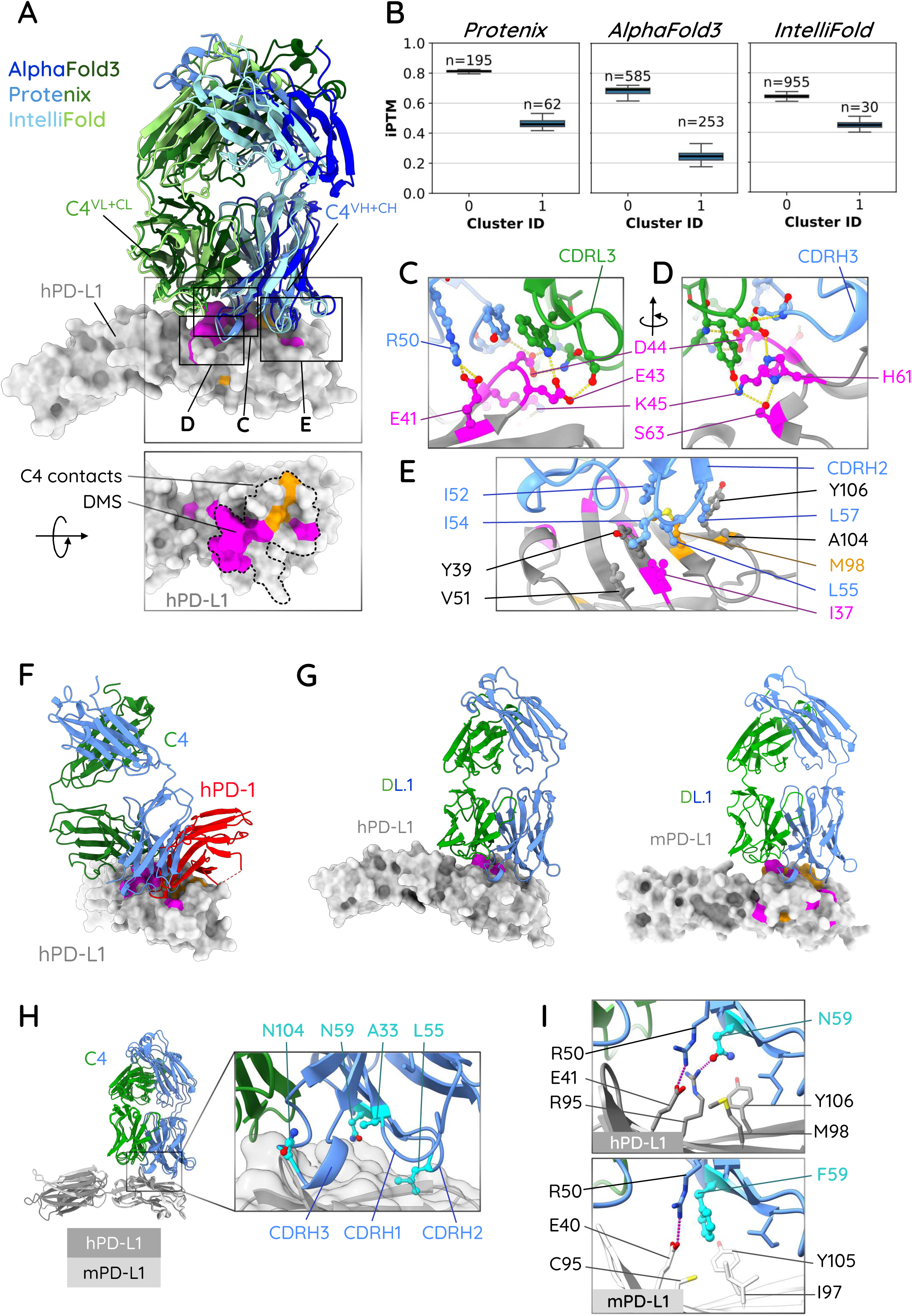
Structural basis of DL.1 recognition of human PD-L1 and comparison of structure predictions. (A) Structural models of the C4 antibody with human PD-L1 (hPD-L1) complex generated by AlphaFold3 (dark blue-green), Protenix (medium blue-green), and IntelliFold (light blue-green), in complex with hPD-L1 (gray surface). The variable and constant domains of the light chain (VL+CL, green) and heavy chain (VH+CH, blue) are indicated. The lower panel highlights the hPD-L1 residues identified by deep mutational scanning (DMS) as contributing to C4 binding (magenta and orange) and residues in contact with C4 antibody as modeled by Protenix are delineated by the dashed line. (B) Distribution of iPTM scores across the predicted binding clusters identified by the three methods. Only clusters comprising more than 50 models and exhibiting iPTM scores > 0.4 are shown. Cluster 0 correspond to the predominant binding mode shown in panel A. (C–E) Close-up views of the predicted C4 - hPD-L1 interface generated with Protenix. Key interactions involve residues from the C4 heavy chain (C) and light chain (D, E) contacting hPD-L1. Residues participating in the interface are shown as sticks and labeled. Predicted contacts include interactions involving hPD-L1 residues E41, D44, K45, and S63, as well as hydrophobic contacts centered on residues I37, V51, I52, I54, L55, L57, M98, A104, and Y106. (F) Superimposition between the model of the C4 antibody–hPD-L1 complex with the experimental structure of the hPD1–hPD-L1 (PDB:) (G) Predicted binding of C4 DLb to human PD-L1 (left) and mouse PD-L1 (mPD-L1, right) showing that the overall predicted binding mode is not changed (H) Structural comparison of the C4 interface on hPD-L1 and mPD-L1. (I) Enlarged views highlight species-specific substitutions at position 59, where N59 in hPD-L1 and F59 in mPD-L1 generate distinct interaction networks with C4, potentially contributing to differences in binding affinity and specificity. The three other substitutions are shown in Supp Fig. 9.

### New sequences generated by Deep Learning are all highly effective with only 4 to 5 mutations

Antibodies generated by our Deep Learning approach were reformatted into IgG for BLI characterization where they exhibited significantly improved affinities compared with parental C4 for human and mouse PD-L1 (Fig. 3D). Their sequences revealed a clear constant motif with subtle variations, providing insight into the contribution of each substitution to affinity (Fig. 3C and Sup. Fig. 5). While A33G and N59F are consistently found across all clones, suggesting a central role in affinity maturation, other substitutions such as S31R or T28D, T28E or G27Y seem interchangeable or dispensable, which might indicate a secondary role. N104A or N104R seem very decisive (Sup. Fig. 5), yet in theory they could be compensated for by other substitutions according to the model. Interestingly, the fourth mutation expected in the quadruplet can be either L55V or G56P, with the absence of one being compensated by the presence of the other. The simultaneous presence of both substitutions does not appear to be required. However, incorporation of only three mutations (A33G, N59F, and N104A) yields a lower affinity than the lowest K_D_ measured on our 10 DMS-only high affinity cross-reactive antibodies. Most likely due to a slightly higher off-rate (Sup. Fig. 6). From this, we infer that L55V or G56P are necessary but less impactful on affinity. These co-dependencies highlight the complexity of the sequence-affinity landscape and underscore the value of the sampling and deep learning strategy used here to explore mutation combinations rather than individual substitutions alone. Overall, the Deep Learning generated variants (DL.1 … 8) clustered tightly in the iso-affinity plot for both human and murine K_D_ values (Fig. 3E), closely mirroring the behavior of the DMS-derived (DMS.1 … 10) variants and demonstrating the successful achievement of high-affinity cross-reactivity toward both human and murine PD-L1. Moreover, all tested clones affinity predictions were consistent with BLI data and were accurately captured by the ensemble model, with “-” corresponding to affinities >100 nM, “+” to nanomolar affinities, and “++” to sub-nanomolar affinities (Sup. Fig. 7).

**Figure 6:**
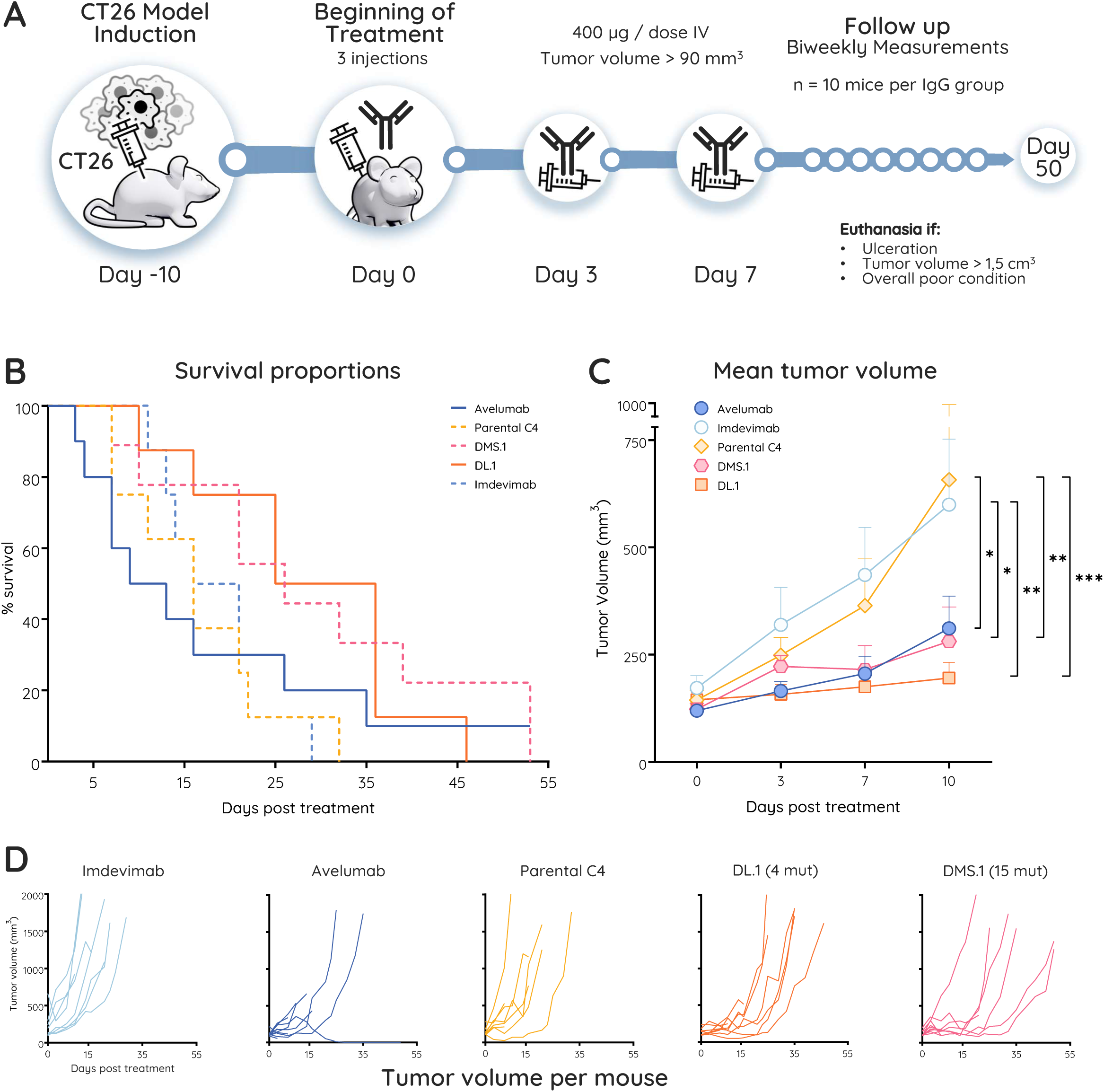
Therapeutic effect evaluation of the engineered antibodies in a mouse colorectal cancer model. (A) Outline of the study. CT26 colorectal cells were implanted subcutaneously in mice and treatment started when sufficient tumor volume of 90mm^3^ was reached. 3 consecutive antibody injections occurred at days 0, 3 and 7. Follow up lasted to day 50. (B) Kaplan-Meier survival curves following the first treatment at day 0. (C) Mean tumor volume per antibody group up to day 10 (first day of euthanasia for reaching maximal 1500mm^3^ volume). Data represent mean ± SEM. A Mixed-effect statistical analysis was performed, * p<0.05, ** p<0.01, *** p<0.001. (D) Details of the individual tumor volume per mouse up till the end of the study (euthanizia).

To verify that the affinity gains observed *in vitro* translate into enhanced recognition of murine PD-L1 expressing cells, binding of both DMS and DL antibodies was assessed on CT26 mouse colorectal cancer cell lines in a FACS saturation assay (Fig. 3F). This murine cell line is known for its relatively high basal expression of PD-L1 [33,34], and is considered as partially responsive to anti-PD-L1 therapies [35,36]. Avelumab, an anti-PD-L1 antibody, and Imdevimab, a non-relevant anti-SARS-CoV-2 RBD antibody, were included as positive and negative controls, respectively (Ill light chain IgG). Apparent EC_50_ from this assay do not reflect affinity because of the multivalent nature of the IgGs and the CT26 cells. Nevertheless, they are consistent with BLI affinities, in that the parental C4 EC_50_ is markedly different from those of all optimized variants. The EC_50_ recorded ranged from 0.18 to 1.02 nM for DMS variants, whereas they ranged from 0.23 and 0.58 nM for the DL variants. Variations are likely attributable to the experimental design. No engineered candidate clearly outperformed the other but they were all considerably improved over parental C4 and the Avelumab control.

### Non-Conserved residues engaged in the epitope of parental and engineered antibodies

To better understand the nature of the substitutions brought during our engineering of selectivity, we set out to map the residues necessary for antibody recognition of the antigen – the functional epitope – by performing the DMS process on the antigen sequence (Fig. 4A). To this end, the membrane distal Ig-like V-type domain from either human or murine PD-L1 was expressed on the surface of yeast cells. Single-mutant libraries were generated to map epitope residues by assessing the binding of monovalent Fab fragments from the parental C4 and two high-affinity cross-reactive variants: DMS.1 generated by DMS with 15 substitutions, DL.1 generated by Deep Learning with four mutations. (Fig. 4A and Sup. Fig. 8). This analysis showed that the functional epitope recognized by parental C4 on human PD-L1 involves residues that are not conserved in murine PD-L1 (Fig. 4B, stripped positions), most likely explaining the poor binding to the mouse antigen.

Affinity maturation altered the binding modalities of the antibodies to human PD-L1 (Fig. 4C). The functional epitope remained centered on the same region, suggesting that the overall antibody-antigen binding mode was preserved. However, local variations were observed with fewer reactive positions detected for the engineered Fabs than for parental C4, and these positions were preferentially conserved between human and mouse PD-L1, as expected for antibodies selected for cross-species recognition. A similar trend was observed on mouse PD-L1, where the engineered Fabs engaged additional positions, particularly among residues conserved between both species (Fig. 4D).

The two high-affinity cross-reactive antibodies displayed largely conserved binding modes toward both human and murine PD-L1, with subtle differences in residue engagement. Epitope of DMS.1 additionally includes H61 for the human antigen (Fig. 4C). The epitope on the murine antigen for DMS.1 involves also Y39, while DL.1 preferentially engages E43, D105, and Y106. Overall, both antibodies targeted the same epitope, but affinity maturation shifted key anchoring interactions toward conserved residues while establishing additional contacts with murine-specific residues, thereby enabling cross-reactive binding.

We then sought to determine whether the C4 antibody was capable of antagonizing the interaction between PD-L1 and PD-1. Overlay of the crystallographic epitope of human PD-1 (Fig. 4E) and the functional epitope of C4 and DL.1 antibodies on human PD-L1 reveal overlapping residues (stripped positions). This suggests that both antibodies likely block PD-1 interaction with PD-L1 similarly to what is observed for Avelumab (Fig. 4F). To confirm the inhibitory capacity of the antibodies against the PD-1/PD-L1 interaction, a competitive ELISA assay was performed (Fig. 4G). This demonstrated that parental C4, DL.1 and SM.1 all compete with human PD-1.

### Docking model generated with a deep-sampling strategy of folding models concur with biological epitope data

To gain deeper insight into the potential mode of interaction between C4 antibody and PD-L1, we used three structure-prediction methods showing the highest success rates for antibody-antigen interactions, AlphaFold3 [27], Protenix-v1 [37] and IntelliFold [38]. We applied these co-folding strategies to sample 1,000 structural models of the human PD-L1 / parental C4 complex, which were subsequently clustered according to their structural similarity. The highest confidence models obtained with all three methods showed strong consistency (Fig. 5A, 5B, Sup. Fig. 9), with a mean interface RMSD (iRMSD) of 0.8805 Å across the three models. The confidence of these interfaces reported by the iptm score between PD-L1 and the heavy and light chains, ranged from 0.67 with IntelliFold to 0.82 with Protenix and no alternative assembly reached such high values (Fig. 5B). Among the three methods, only a local difference in the orientation of the tip of the CDRH3 loop conformation was observed in the AlphaFold3 model (this variation is discussed in N104 mutation analysis below). Interestingly, all three methods generated models consistent with the DMS-based epitope mapping shown in the lower panel of Fig. 5A, although this information was not used to guide the docking. A buried salt-bridge between R50^C4^ and E41^hPD-L1^ is observed at the center of the interface (Fig. 5C). This interaction is likely a major determinant of the complex stability and provides a plausible structural explanation for the strong sensitivity of the substitutions at position 41 revealed by DMS. Adjacent to E41, a dense cluster of charged and polar residues in human PD-L1 (Fig. 5C, D) forms a hydrogen-bonding network with side-chain and backbone atoms of C4 residues predominantly located in the CDRL3 loop. A second major anchoring region (Fig. 5E) involves hydrophobic residues I52, I54, L55 and I57 in CDRH2 of the C4 antibody in its CDRH2 and is predicted to lie within the apolar patch of PD-L1 formed by I37, Y39, V51, M98, A104 and Y106 (Fig. 5E). This apolar patch is also involved in the interaction with PD-1, consistent with the experimental observation that C4 and PD-1 compete for binding to PD-L1 (Fig. 5F).

Overall, the conformational convergence of the different models toward the same relative orientation between the mutated antibodies and the two PD-L1 orthologs further supports the conclusion drawn from the DMS experiments that the complex structures were not substantially altered by the introduction of the mutations.

Given the convergence and consistency of the structural models with the experimental data, we next explored whether the effects of the minimal set of mutations required to generate a C4 antibody capable of binding both human PD-L1 and mouse PD-L1 could be rationalized. We therefore tried to investigate the contribution of substitutions present in the sequence of DL.1, namely N59F^C4^, L55V^C4^, N104A^C4^ and A33G^C4^ (Fig. 5H). Comparative analysis of the structural models provides a possible explanation for the effect of the N59F ^C4^ substitution (Fig. 5I) while additional model-based interpretations of the other mutations are provided in the Supplementary Data (Sup. Fig. 9). In human PD-L1, the region contacting residue N59 ^C4^ contains an arginine (R96 ^hPD-L1^), which is predicted to form a hydrogen bond between its guanidinium group and the side chain of N59 ^C4^, thereby promoting favorable solvation of the interface. In mouse mPD-L1, this arginine is replaced by a cysteine (C95 ^mPD-L1^), which is considerably less bulky and favors more hydrophobic interactions. The N59F^C4^ substitution creates an apolar cluster involving C4 residue F59 ^C4^ and residues C95 ^mPD-L1^, Y105 ^mPD-L1^, and I97 ^mPD-L1^ of mouse PD-L1. We hypothesize that the N59F ^C4^ substitution is not too detrimental to binding human PD-L1 because a cation-π interaction may be maintained between F59 ^C4^ and R96 ^hPD-L1^.

### Antibodies engineered with minimal mutations by Deep Learning have an *in vivo* therapeutic effect in a CT26 colorectal mouse model of cancer

Together, the epitope-mapping and structural analyses suggested that the engineered antibodies retained a therapeutically relevant binding mode while acquiring improved recognition of murine PD-L1. We therefore sought to determine whether these properties translated into *in vivo* efficacy in a preclinical cancer model.

After 10 days of induction of the subcutaneous CT26 model, mice with tumor volume above 90 mm^3^ were selected and were randomly assigned in 5 groups of 10 mice (Fig 6A). At days 0, 3 and 7, the groups received an intravenous injection of a therapeutic dose of their respective antibodies (Avelumab, Parental C4, DMS.1, DL.1 or Imdevimab as an isotypic control). Overall, only 1 mouse survived this harsh study, in the Avelumab group (Fig. 6B).

Survival analysis show a clear divide between the engineered antibodies and the parental antibody which is indistinguishable from the isotypic control (Fig. 6B). Engineered antibodies clearly prolonged survival compared to control groups (median survival of 26- and 30.5-days vs 11 to 18.5 days for engineered and control groups, respectively). The Avelumab group suffered from ulcerations right after treatment, resulting in in early euthanasia (before day 10 post treatment), and therefore the proper therapeutic efficacy of Avelumab could not be observed during the first 10 days. After day 15, the descent of the curve slowed down and became more representative of the expected effect of Avelumab. These tendencies were also reflected in the mean tumor volume of each group (Fig. 6C) compared to the control Imdevimab and parental C4 groups, which is significant at 10 days post treatment. At this point, the parental C4 and Imdevimab had similar effects, whereas the 3 other antibodies similarly contributed to keeping mean tumor volume low. Individual tumor volumes (Fig. 6D) again showed the therapeutic effect of Avelumab and our engineered antibodies on tumor growth, compared to Imdevimab and parental C4, on delaying the exponential growth of CT26 subcutaneous tumors. These data clearly show that the parental C4 antibody exhibits low therapeutic effect in mice, with minimal difference from the negative control. In contrast, the optimized versions, DMS.1 and DL.1, demonstrate similar efficacy to Avelumab. These results further highlight that the minimal mutations detected by our Deep Learning approach were sufficient to generate effective therapeutic anti PD-L1 antibody candidates with cross-reactivity for both mouse and human PD-L1.

## Discussion

### Combining DMS and Machine Learning for cross-reactivity engineering

Throughout this study, we aimed to develop an affinity maturation strategy that could reach the best possible compromise between top-tier affinity and low mutation count. First, we devised a combinatorial library of engineered antibodies and screened it to recover high-performing ligands displaying equivalent affinity for both human and mouse PD-L1. This approach successfully yielded ten cross-reactive variants of C4, one of which, DMS.1, was fully characterized as a therapeutically relevant immune checkpoint blocker in mouse cancer models. Subsequently, we aimed to uncover the precise effect of each mutation so as to engineer candidates with the highest affinity using the minimal set of mutations possible. For this, we leveraged Deep Learning trained on DMS-derived sequence-affinity data to identify minimally mutated variants predicted to retain high-affinity cross-reactivity. The resulting antibodies, harboring only 4 to 5 mutations while preserving their modes of antigen binding, provide a panel of highly human, cross-reactive candidates suitable for preclinical studies while illustrating the potential of Machine Learning to guide efficient antibody optimization.

### Deconvoluting mutational contribution through Machine Learning

Deciphering the contribution of individual substitutions is challenging, particularly when multiple mutations are recurrently co-selected, making their individual effects difficult to disentangle. Based on expert analysis, ubiquitously retained mutations could not be discerned from one another. Additionally, while visualizing enrichments via sequence logos can reveal conserved positions, but they do not resolve mutation co-dependencies or discriminate passenger mutations from essential ones. Interestingly, the manually designed variants, selected through manual analysis of Sanger sequencing data, were the only candidates that failed to achieve sub-nanomolar affinity for murine PD-L1. Although informed by experimental data, this expert-driven approach appeared less effective than the machine learning strategy at identifying optimal combinations of mutations, underscoring the ability of the model to exploit sequence–function relationships that are not readily apparent from manual inspection alone.

Our ML-based approach helped us make sense of this complex sequence-affinity landscape. Global models outperformed other architectures on this dataset, and a ConvNeXt-based CNN was selected as the preferred model. Using an ensemble-model, we reverse-engineered the least-mutated sequences that still meet our affinity criteria. This approach proved successful as every single one of our predictions was validated *in vitro*.

The definition of the minimal mutational footprint remains intrinsically linked to the performance objectives being targeted. We initially adopted a conservative strategy when selecting deep learning-generated sequences for experimental validation. In hindsight, we are now very confident that – provided we remain within the bounds of the model – any sequence we generate from it will have suitable *in vitro* performance.

This study also contributes to the growing evidence that antibody sequence-function relationships can be learned by machine learning. Previous work established that deep learning architectures can distinguish binders from non-binders [39]. We expand upon this finding by predicting affinity ranges - categorized as low, medium, and strong binders. More recently, related deep mutational learning strategy [30] have been used to predict the resistance of therapeutic anti-SARS-CoV-2 antibodies against ACE2-compatible antigenic drift and to prioritize antibody combinations resilient to viral escape. In contrast, our approach uses DMS-coupled deep learning as an active antibody-engineering framework which does not only evaluate how antibodies withstand antigen variations, but also confer cross-species reactivity with controllable sequence divergence.

Importantly, the combination of DMS and deep learning also generated a rich experimental framework for structural interpretation. When structural models were generated with high confidence and showed full consistency with the epitope-mapping and binding data, they provided plausible explanations for the effects of selected mutations. In particular, some substitutions, such as N59F and L55V, appear to directly affect the antibody-antigen interface, whereas others, such as A33G and N104A, may act more indirectly by modulating the stability of CDR loop conformations. These interpretations remain speculative and will require experimental structural validation. Nevertheless, they illustrate how AI-based structural modeling can synergize with DMS by helping to rationalize mutation effects in a systematic manner, rather than serving solely as a visualization tool.

This perspective is particularly relevant in light of recent structure-informed antibody design approaches, in which structural models and language models have been used to propose beneficial mutations in specific antibody-antigen complexes [40]. Our study differs in that the model was trained on context-specific DMS data and used to identify minimal mutation combinations with experimentally validated cross-species activity. Together, these approaches suggest that future antibody engineering workflows could integrate experimental DMS, sequence-based learning and structural modeling more tightly, with structural models not only rationalizing selected variants but also helping to prioritize substitutions for subsequent experimental sampling.

### Balancing affinity and antibody humanness

Importantly, this approach generates an entire in silico repertoire of candidates, rather than the limited number obtainable through colony picking. This new repertoire of functionally enhanced clones can also be screened for other desirable properties. In this study, we sought to optimize sequence humanness by generating antibodies as close as possible to the corresponding human germline, with the goal of minimizing potential immunogenicity. Although this study focused on generating high-affinity cross-reactive antibodies, the same framework could also be used to fine-tune affinity. This may be relevant for bispecific or multispecific antibody engineering, where the affinity of each binding arm must be fine-tuned according to antigens identity and their cell-surface density [41–43]. It may also be relevant in oncology, where intermediate affinities can sometimes be advantageous as excessively tight binding may impair tumor penetration or promote target-mediated toxicity, whereas overly weak interactions may reduce efficacy [44–47].

The generation of a large in silico repertoire also creates opportunities beyond affinity maturation alone. Instead of being limited to the few candidates obtained by colony picking, this repertoire can be screened for additional desirable properties. In this study, we prioritized sequence humanness with the goal of limiting potential immunogenicity. In addition, complementary filters or predictive models could be integrated into the design workflow in particular in regard to mAb developability. The incorporation of previously described tools [48–50] to filter candidates with developability liabilities could further facilitate the identification of a balanced trade-off between affinity and downstream therapeutic potential. [51]. More broadly, the DMS-coupled deep learning strategy described here could be highly relevant to further engineer antibodies not solely to maximize affinity, but also to generate antibodies with tunable, on-demand binding properties tailored to specific preclinical or therapeutic applications.

## Acknowledgements

This work was supported by state aid managed by the National Research Agency under the France 2030 program, referencing ACCREDIA ANR-22-PEBI-0009.

## Methods

### Design of DMS libraries

For affinity maturation: deep mutational scanning (DMS) libraries of Fab variants with single amino acid mutations were generated by splicing by overlap extension PCR (SOE-PCR) using degenerate NNK primers in oligopool format (IDT). A library was constructed for the four targeted regions: VH positions 1 to 62 and 63 to 122 and VL positions 1 to 56 and 57 to 109. For epitope mapping: PD-L1 expression on yeast was previously tested in clonal Wild Type format. Binding was observed for the full form of PD-L1 and the Nter part. DMS libraries were thus only generated for the Nter part of the sequence taken from either PDB entry 3BIS (residues 1 to 122) or GenBank entry AAG18509.1 residues (18 to 133) for human PD-L1 and mouse PD-L1 respectively.

### Yeast Surface Display of Fab or PD-L1

Preparation of competent yeast cells EBY100 (ATCC® MYA-4941) and library transformation were performed according to Benatuil *et al*. [52] Gap repair transformations were made in a pSWG plasmid by restriction using sites around the zones of interest : Fab VH and VL, and PD-L1 Nter domain. Around 4µg of SOE-PCR generated insert DNA were mixed with 1-2 µg of digested backbone for an efficiency of around 10^7^ colony forming units (cfu) per electroporation. After transformation, cells were cultivated in 250 mL of SD-CAA medium [53] (1.7Illg/L yeast nitrogen base without casamino acids and ammonium sulfate, 5 g Ammonium sulfate, 20Illg/L glucose, 5Illg/L casamino acids, 100IllmM sodium phosphate, pH 6.0). After a passage to an OD_600_ of 0.25, cells were grown at 30°C until OD_600_ 0.5–1.0 and re-suspended in the minimal volume of SG-CAA for induction, so as to conserve 10 times the number of cfu obtained per transformation. Induction time varied from 18h to 72h with no discerning impact on yeast Fab presentation (>70% of cells induced is ideal).

### Flow cytometry

For library sorting, 10^7^ to 2.10^8^ induced cells were washed in PBSF (phosphate-buffered saline (PBS), bovine serum albumin (BSA) 0.1%) and resuspended in PBSF containing the appropriate antigen concentration (50010-M08H-B and 10084-H08H-B for mouse and human biotinylated CD274 respectively, Sinobiological) using an appropriate volume to avoid ligand depletion as performed in Hunter et al. [54]. After 2 hours of incubation at 20°C with agitation, cells were washed with ice-cold PBSF to avoid dissociation. Cells were incubated on ice for 25 minutes with anti-HA antibody (Invitrogen HA Tag Mouse anti-Tag, DyLight® 650 conjugate, Clone: 2–2.2.14; 1:100 dilution) and Streptavidin-PE (Thermo Fisher Scientific; catalog number S866; 1:200 dilution). Cells were subsequently washed with ice-cold PBSF and sorted with a BD FACS Aria™ III cytometer using BD FACSdiva™ software.

### Deep sequencing and analysis of NGS data for DMS

Plasmid DNA of each yeast population was extracted using Zymoprep Yeast Plasmid Miniprep II and prepared for sequencing as described in Medina-Cucurella and Whitehead [55]. Two-step PCR was performed to amplify the region of interest and add Illumina adapters and barcodes for multiplexing. Deep sequencing was performed with an Illumina ISeq 100 device (2x150 bp, 300 cycles) with at least 200,000 reads per population. Reads were demultiplexed, and each sample was processed separately using a custom R script. Paired end reads were joined with NGmerge. DNA sequences were translated in protein sequences and identical sequences were grouped. Sequences not repeated at least two times were filtered out. Single-mutants were selected to allow calculation of enrichment ratios for each single mutation.

### Library generation et recombination

Candidate libraries combining the positions identified by DMS were compiled using degenerate primers from Eurofins. Codon libraries were designed using SwiftLib (http://rosettadesign.med.unc.edu/SwiftLib/). Libraries were cut in half accros the Variable Heavy domain of the Fab so as to maintain a diversity that was manageable for yeast transformations. After rounds of sorting, sub-libraries were pelleted and extracted with Zymoprep. Through PCR, the two sub-libraries were joined as one by simple hybridation in a constant part in the middle. Gap-repair transformations were performed as prior, while aiming to have 10 times more cfu than the theoretical codon diversity of each library.

### Machine Learning Loops

Dataset building and learning loops were developed following the pytorch documentation (https://docs.pytorch.org/tutorials) and through scikit-learn packages. NGS data was demultiplexed, QC performed with R package dada2, low quality and rare sequences were discarded. Common sequences across classes as well. Amino acid sequences were one-hot encoded using scikit-learn packages. Sequences of 122 amino acids are converted to matrices of 122 by 20 (for each amino acid) with a 1 or a 0. This vector is associated with a category encoded by 0, 1 or 2. Train / Test ratio was constant at 25 % for model type comparison and cross-validation. But some leeway was given when generating the final models for prediction of sequences 82/18%) while empirically searching for the best performance. Hyper parameter search was done empirically. Model learning was stopped at the best performing epoch when test metrics were still superior to the train ones. Sequences to predict were generated in R by reverting substitutions from the Wild Type to sequences obtained by saturation mutagenesis prior. All combinations were fed to the 3 best performing models. Final sequence selection for *in vitro* testing was performed by looking for high confidence predictions across all 3 models.

### Production and purification of IgGs

All IgG Wild Type and variants were obtained by transient transfection of ExpiCHO™ (Thermo-Fisher) at a density of 2.5 10^6^ cells/mL in 10-400 mL medium using the dedicated ExpiFectamine™ kit and reagents and about 0.8µg of heavy chain plasmid and light chain plasmid each per mL of transfected cell. Synthetic genes corresponding to selected VH mutants were generated by template free PCR as described by Xiong et al., [56] using primers designed by Mutation Maker [57] (https://github.com/Merck/Mutation_Maker). They were cloned into the mammalian expression vector pcDNA 3.4 with a CMV promoter and all the same CH1-CH2-CH3 gene or C_ll_. After transient expression, centrifuged and filtered supernatants were purified using HiTrap Protein G for all variants, following the manufacturer’s instructions (GE Healthcare). Size-exclusion chromatography was performed (HiPrep Sephacryl-S-200 HR or S100HR) with PBS.

### Size-exclusion chromatography

A Superdex-200 10/300 GL (GE Healthcare) column was used on an AKTA Pure system (Cytiva). Runs were monitored with Unicorn TM 7.6 software (Cytiva). Sample was applied on the column using a flow rate of 0.5 mL/min with PBS as running buffer.

### Affinity measurement by BLI

Binding kinetics were determined using an Octet RED96 instrument (ForteBio). Anti-hIgG Fc Capture (AHC) Biosensors (Fortebio) were loaded with IgG molecules (100 nM) for 60 seconds. After baseline using kinetic buffer (PBS, BSA 0.5% (w/v) and Tween 20 0.05% (v/v)), association of un-tagged murine or human PD-L1 (50010-M08H, 10084-H04H, Sinobiological) was measured at different concentrations (50 nM to 0.78 nM) for 300 seconds before dissociation in kinetic buffer for 1500 seconds. For the wild type IgG, prior to affinity maturation, a fit was only satisfactory using a short association time of 100s and higher concentrations of antigen (150 nM to 2.3 nM murine PD-L1). Analysis: Data of the control without antigen were subtracted from all binding curves, and binding kinetics were fitted using a global 1:1 Langmuir-binding model. Iso affinity graphs were plotted with GraphPad prism 8.

### ELISA

PD-1 / PD-L1 inhibition screening performed with ELISA kit EP-101 from Acrobiosystems following manufacturer guidelines and protocols. Control and Test antibodies were used at the same starting concentration as the kit reference anti-PD1-neutralizing antibody: 10µg/mL and diluted by 2 in series to reach 0.078 µg/mL as the lowest concentration. Background signal removal, normalization, analysis and plotting was performed using GraphPad Prism 8.

### Docking model generation

Structural modeling was performed using three different AI-based methods which were benchmarked on antibody-antigen complexes with similar performance, namely AlphaFold3 [27], Protenix-v1 [37] IntelliFold [38]. For each of these methods, 1000 models were generated per run, using 200 seeds selected at random.

### Structural-clustering algorithm

To reduce the redundancy of the generated models, a structural clustering was applied per run. DockQ [58] (https://github.com/wallnerlab/DockQ) metric was used to compare the generated models in a pairwise manner. The interface confidence score ipTM restricted to the antibody-antigen interface of each structural model was used to assess prediction quality likelihood. Taking into account the antibody heavy-light chains interface led to over-confident docking estimations. The antibody-antigen ipTM was obtained by parsing the JSON confidence files and averaging matching pairwise ipTM values. We performed an iterative clustering, selecting the cluster center according to the maximal value of the antibody-antigen ipTM, and then gathered similar generated-structures using DockQ. A threshold of 0.23 was used to form clusters, to obtain structures with an “Acceptable” DockQ classification inside a same cluster. The method converges when all structures have been assigned a cluster id. Cluster representatives are models with the highest confidence (ipTM) per cluster. The selected models are available in ModelArchive at https://www.modelarchive.org/doi/10.5452/ma-2lydk.

### Cell line

Murine colon adenocarcinoma CT26 cell line CRL-2638 ATCC (ATCC) was cultured in DMEM supplemented with 10% of fetal bovine serum and 1% antibiotic.

### *in vitro* cell binding assay

500 000 CT26 cells were resuspended in RPMI without FBS. For the blocking conditions, cells are first incubated with each unconjugated anti-PD-L1 antibody (ratio 1:100 - conjugated: unconjugated antibody) at 37°C for 1hour. Then, for both saturation and blocking conditions, cells were incubated with each biotin-conjugated anti-PD-L1 antibody, at 4°C for 4 hours. Cells were then washed with PBS and centrifugated (500g, 5min) 3 times, before incubation with streptavidin-PE (Invitrogen) at 4µg/mL at room temperature for 30 minutes. Cells were then washed with PBS and centrifugated (500g, 5min) 3 times before reading with Attune NTX flow cytometer (ThermoFisher).

### *in vivo* treatment

Healthy female Balb/c mice, 6-weeks of age (Janvier Lab), were subcutaneously implanted into one flank with 1.10^6^ CT26 cells in 100µL of PBS. Treatments were initiated once tumors reached 90mm^3^, corresponding to D0. Mice received 3 intraveinous injections at D0, D3 and D7, of 400µg of antibody per dose, for a total dose of 1.2mg of antibody. Mice were randomly divided into 5 groups, receiving either avelumab (N=10), imdevimab (N=8), C4 WT (N=8), C4 DMS (N=9) or C4 IA (N=8). Tumors’ volumes were assessed twice per week and estimated according to the formula: 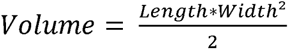. Euthanasia was performed when maximal volume of 1500mm^3^ was reached, or became ulcerated. Mice with ulcerated tumors before D10 were excluded (N=4, N=2 and N=1 in the avelumab group C4 WT and C4 DMS groups, respectively).

### Statistical analysis, graphs and schematics

All Illustrations were made with Blender 4.5 LTS; heatmap were plotted with R; sequence logos plotted with R package ggseqlogo and colored in paint.Net. Iso-affinity plots were made with Prism. GraphPad prism software (v11.0) was used for *in vivo* statistical analysis. A two-way ANOVA with a Tukey post-test was performed to evaluate statistical differences between the tumor volumes in all 5 groups between day 0 and 10. A logrank test was performed to analyze survival data. Statistical significance was set-up at p<0.05.

## Supplementary Figures Legends

**Supplementary figure 1: DMS heatmaps for Variable Heavy chain** Enrichment analysis of a library of single mutants on the VH of the parental antibody in various conditions. (A) positive gate (higher affinity) on mouse PD-L1, (B) Negative gate (Binding Lost) on mouse PD-L1, (C) positive and (D) negative gate on human PD-L1. (E) Final heatmap after concatenation of all criteria in fig1A. (F) Combinatorial library built with degenerate codons with Switftlib (see methods). Yellow: amino acids to include from panel E. Green: Parental amino acid to introduce to the library. Red: unwanted amino acids added as a compromise from using degenerate codons.

**Supplementary figure 2: DMS heatmaps for Variable Light chain** Enrichment analysis of a library of single mutants on the VL of the parental antibody in various conditions. (A) positive gate (higher affinity) on mouse PD-L1 or (B) on human PD-L1. (C) Final heatmap after concatenation of all criteria in fig1A. (D) Combinatorial library built with degenerate codons with Switftlib (see methods). Yellow: amino acids to include from panel C. Green: Parental amino acid to introduce to the library. Red: unwanted amino acids added as a compromise from using degenerate codons.

**Supplementary figure 3: Sequences and BLI graphs of Antibodies engineered by DMS and derivatives** (A) Sanger sequencing of all colony-picked clones. Consensus amino acids in comparison with the parental sequence are presented as a dot. Below the dotted line (SM.11 to SM.16) are sequences added in the logotype image that have not been expressed as IgG nor tested in BLI, though they were recovered in the same colony picking method as SM.1 to SM.10. (B) Humanity content of parental C4 and DMS engineered sequences. Obtained with IGMT Gap-Align (imgt.org/IMGTindex/IMGTDomainGapAlign.php) when compared to IGHV1-69 allele 04 from *Homo sapiens*. (C) BLI graphs used to plot iso affinity in figure 1. Either on human (orange) or mouse PD-L1 (blue). For SM.1: tip 1 (50nM) excluded from analysis. (D) Sequences generated by iteratively adding substitutions with high enrichment in the logo of colony picked candidates. Only amino acids differing from the parental C4 are shown. These were produced as IgG by transient expression of mammalian cells. (E) BLI affinity assessment of IgGs from (D).

**Supplementary figure 4: Learning curves of the components of the ensemble model** Train / Test loss and accuracy per epoch of learning on each dataset A, B or C as presented in figure 2. Black arrow denotes the epoch at wich the model was saved for use in predictions.

**Supplementary figure 5: Every 4 mutation sequences voted by the ensemble models as “++”** (A) Sequence comparison of these 9 sequences against the parental C4 (first row). Consensus amino acids are represented as a dot. (B) Details of the predictions by the ensemble model with predictions of each single model and the confidence metrics given by the model when deciding. A color code was added on the confidence metric for class 2 (“++”): red < 80%, green > 90%, orange in between.

**Supplementary figure 6: Analysis of 3 mutations sequences** (A) All sequences harboring 3 mutations retained as class 1 (“+”) by the ensemble model. Comparison against the parental C4 (first row). Consensus amino acids are represented as a dot. (B) Details of the predictions by the ensemble model with predictions of each single model and the confidence metrics given by the model when deciding. A color code was added on the confidence metric for class 2 (“++”): red < 80%, green > 90%, orange in between. (C) BLI affinity assessment of sequence 3.E as an IgG, compared to variants described in the main figures, representing the desired affinity.

**Supplementary figure 7: Every experimental K_D_ value compared to model predictions of antibody behavior** Rightmost row (experimental K_D_ values) are color coded according to the scale on the right. Starting from the left, columns 3, 4, 6 and 8 are colored with a discrete code: 0 “-“ → orange ; 1 “+” → green ; 2 “++” → blue.

**Supplementary figure 8: DMS on PD-L1 for epitope mapping heatmaps** (A) All unmodified heatmaps obtained for the 5 epitope mappings. Coloring in white and pink for values above 0. Values in log of enrichment. Below each heatmap, residues considered as a part of the epitope are colored in pink if over 15 amino acid substitutions in the column above reach the cutoff (significant enrichment). Residues mildly engaged are shown in orange if 10-14 amino acid substitutions in the column above reach the cutoff. (B) Solvent Accessible Surface Area obtained with pymol. Residues that meet the criteria for being part of the epitope but have a SASA < 10% are considered buried and excluded from the epitope. (C) Threshold detection for the heatmaps: Each box is a plot of all the values of each heatmap. Revealing the difference between medium enrichment and significant enrichment. Values are normalized per heatmap between 0 and 1 and threshold is set to arithmetic mean + standard deviation. (D) Models from figure 4 with the epitopes colored

**Supplementary Figure 9: Structural analysis of species-specific recognition determinants in the C4-PD-L1 interface** (A) Schematic overview of the structural prediction and analysis pipeline used to generate and evaluate antibody / PD-L1 complex models. Three methods, namely AlphaFold3, Protenix-v1 and IntelliFold have been used. 1000 structural decoys generated with every folding method are clustered iteratively using DockQ structural comparison. (B) Structural role of residue L55 in C4 and predicted effect of the V55 substitution. The residue is positioned at the center of the binding interface and establishes contacts in both hPD-L1 and mPD-L1 models. Comparison of C4 and C4 DL.1 models highlights the conservation of the overall binding mode with local rearrangements in the vicinity of L55/V55. The effect of the L55V substitution is directly linked to the importance of the hydrophobic patch at the surface of both human and mouse PD-L1 for the interaction with C4. We hypothesize that removal of one methyl group from the side chain at position 55 enables more optimal packing of aromatic and aliphatic side chains involved in the apolar interaction at this interface. The lower panel shows the superposition of the hPD-L1- and mPD-L1-bound models, illustrating the subtle differences in side-chain packing that could be brought by the mutation. (C) Structural consequences of substitutions at position 33. Although substitutions to glycine are generally thought to be unfavorable because of the increased flexibility and conformational freedom they confer, it is not the case for residue 33, that is predicted to be part of a β-strand. In the human PD-L1 complex, residue A33 is associated with the formation of a small cavity that can accommodate only a single predicted interfacial water molecule. In contrast, substitution to G33, which is favored in both hPD-L1 and mPD-L1 according to DMS data, enlarges the cavity and is predicted to stabilize an extended water-mediated hydrogen-bonding network. Deep mutational scanning (DMS) data for position 33 are shown alongside the structural models and support the enrichment of residues compatible with water-mediated interactions at this site. This model also provides a plausible explanation for the specific DMS pattern observed at position 33: the side chain of Q33, which contains both hydrogen-bond donor and acceptor groups at its terminus, could occupy the cavity and establish a hydrogen-bonding network with S35 side-chain and L106 peptide backbone carboxy group, analogous to that predicted for the interfacial water molecules. (D) Structural comparison of models containing residue 104 substitutions. AlphaFold3 and Protenix predict distinct conformations of the interface surrounding residue 104, leading to different packing arrangements between the antibody and PD-L1. In the AlphaFold3 model, the CDRH3 loop adopts an extended conformation, in which N104 points toward the solvent and does not make any contact at the interface. In the AlphaFold3 models, CDRH3 is folding upon A104 mutation, allowing its apolar moiety to interact with the Y50-L56-A108 apolar pocket and allowing the formation of the β-turn as predicted by Protenix. This conformation could favor a higher packing at the interface between C4 and PD-L1. This interpretation would be fully compatible with the spectrum of effects observed in the DMS of position 104. Notably, the most polar substitutions (in Q, D, N or E) at this position were deleterious, whereas substitutions to hydrophobic residues or residues containing large aliphatic moieties were generally favorable. This pattern is consistent with a mechanism in which sidechains with apolar moieties promote packing within the apolar pocket at the VH-VL interface, stabilize the β-turn conformation at the tip of CDRH3 loop and thereby favor formation of a more compact and stable interface with PD-L1. (E) Comparative structural predictions of AlphaFold3 and Protenix obtained for all combinations of C4 variants and human (hPD-L1) or mouse (mPD-L1) PD-L1 orthologs, except for wild type C4 – human PD-L1 complex show in Figure5. Left: C4–PD-L1 couple modeled. Middle: distribution of iPTM scores across the predicted binding clusters identified by AlphaFold3 and Protenix. Only the two best clusters (if there are two) are represented. The selected cluster is indicated by a red contouring. Right: predicted binding mode of the selected cluster and associated iPTM score, showing that the overall confidently predicted binding mode is not changed. No docking solution similar to the reference binding mode has been found for C4 wild type–Mouse PD-L1: all generated clusters have a low interface confidence which might indicate that both folding models are sensitive to the poor affinity between the two partners.

