## Supplementary figures for "Machine Learning-Guided Engineering of High-Affinity Cross-Reactive Antibodies with Minimal Mutations"

Supplementary figure 1: Variable Heavy chain Heatmaps and library design

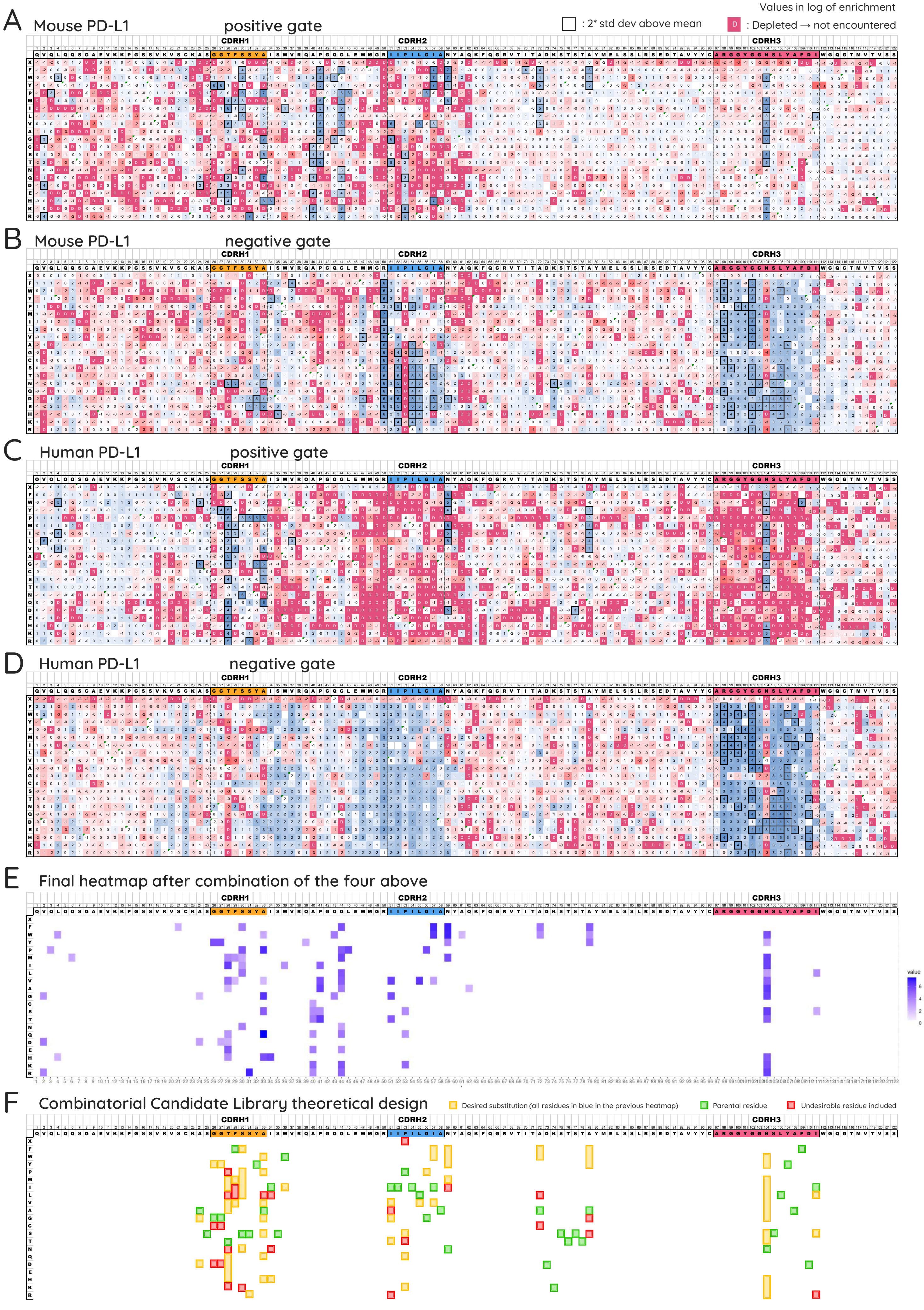

**Supplementary figure 2:** Variable Light chain Heatmaps and library design

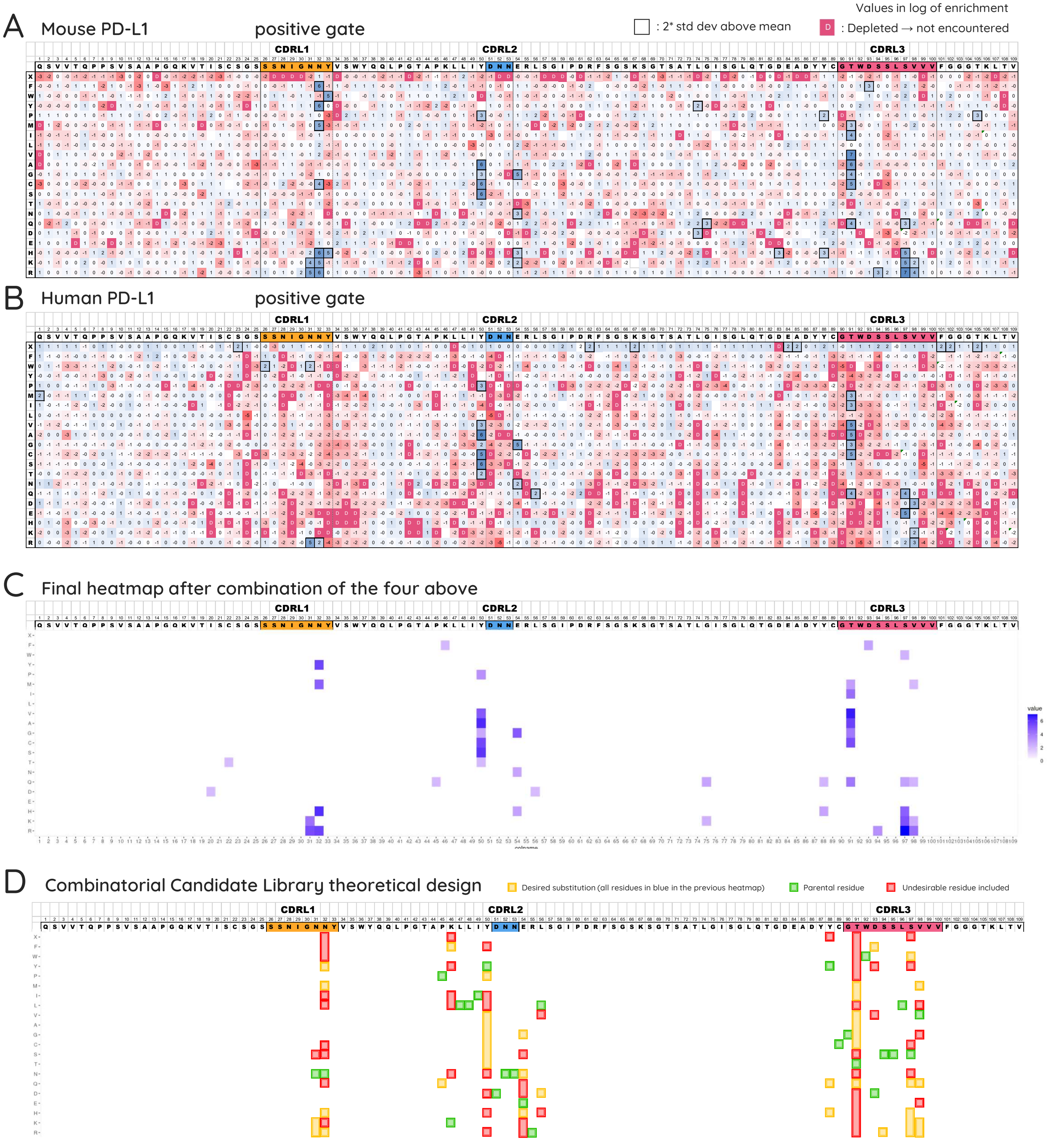

Supplementary figure 3: Sequences and BLI graphs of Antibodies engineered by DMS and derivatives

A

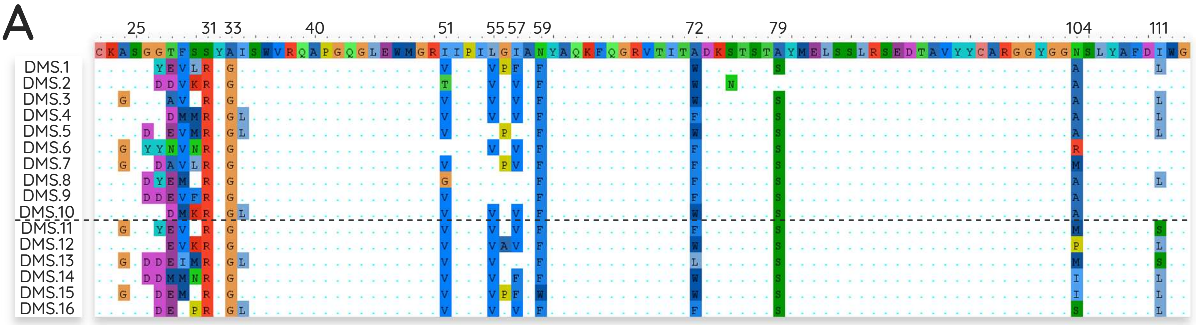


B

| | % Identity |
|---|---|
| Parental C4 | 99 |
| DMS.1 | 85,7 |
| DMS.2 | 86,7 |
| DMS.3 | 87,8 |
| DMS.4 | 86,7 |
| DMS.5 | 86,7 |
| DMS.6 | 85,7 |
| DMS.7 | 85,7 |
| DMS.8 | 88,8 |
| DMS.9 | 87,8 |
| DMS.10 | 86,7 |

*H. Sapiens*

IGHV1-69 *04

C

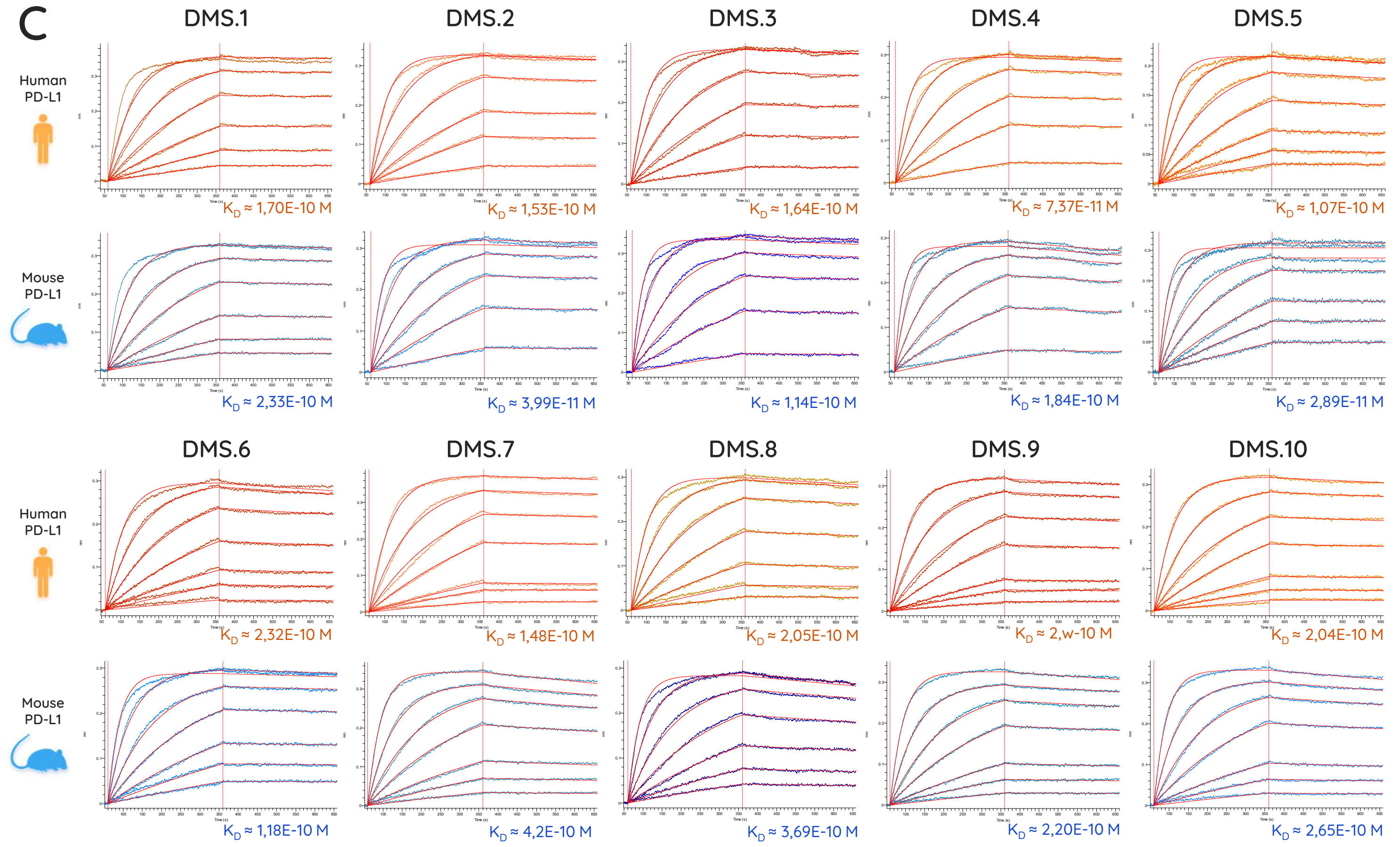


D

### Hand-made minimal mutations sequences

| 24–36 | 50–59 | 71–79 | 97–111 | Mutations |
|---|---|---|---|---|
| ASGGTFSSYAISW | RIIPILGIAN | TADKSTSTA | ARGGYGGNSLYAFDI | |
| .......R..... | .......... | ......... | ............... | 1 |
| .......R.G... | .......... | ......... | ............... | 2 |
| .......R.G... | .........F | ......... | ............... | 3 |
| .......R.G... | .V...V...F | ......... | ............... | 5 |
| .......R.G... | .V...V...F | .F....... | ............... | 6 |
| .......R.G... | .V...V...F | .F....... | .......A....... | 7 |

* 50 nM tip discarded

E

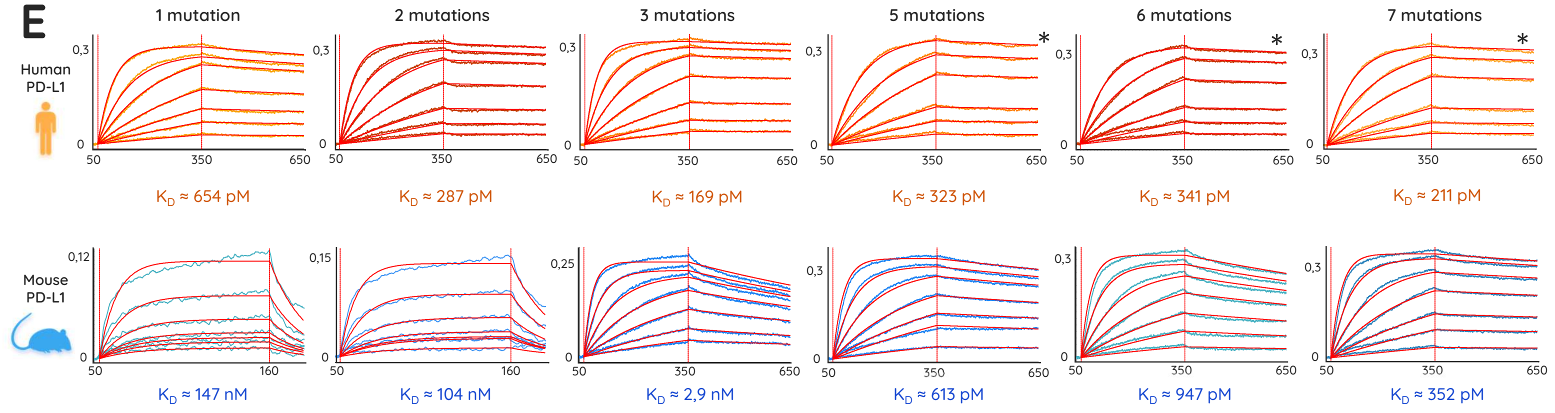

Supplementary figure 4: Learning curves of the components of the ensemble model

**Model A** (on dataset A) — Epoch 9, Acc = 0,905

**Model B** (on dataset B) — Epoch 10, Acc = 0,92

**Model C** (on dataset C) — Epoch 5, Acc = 0,875

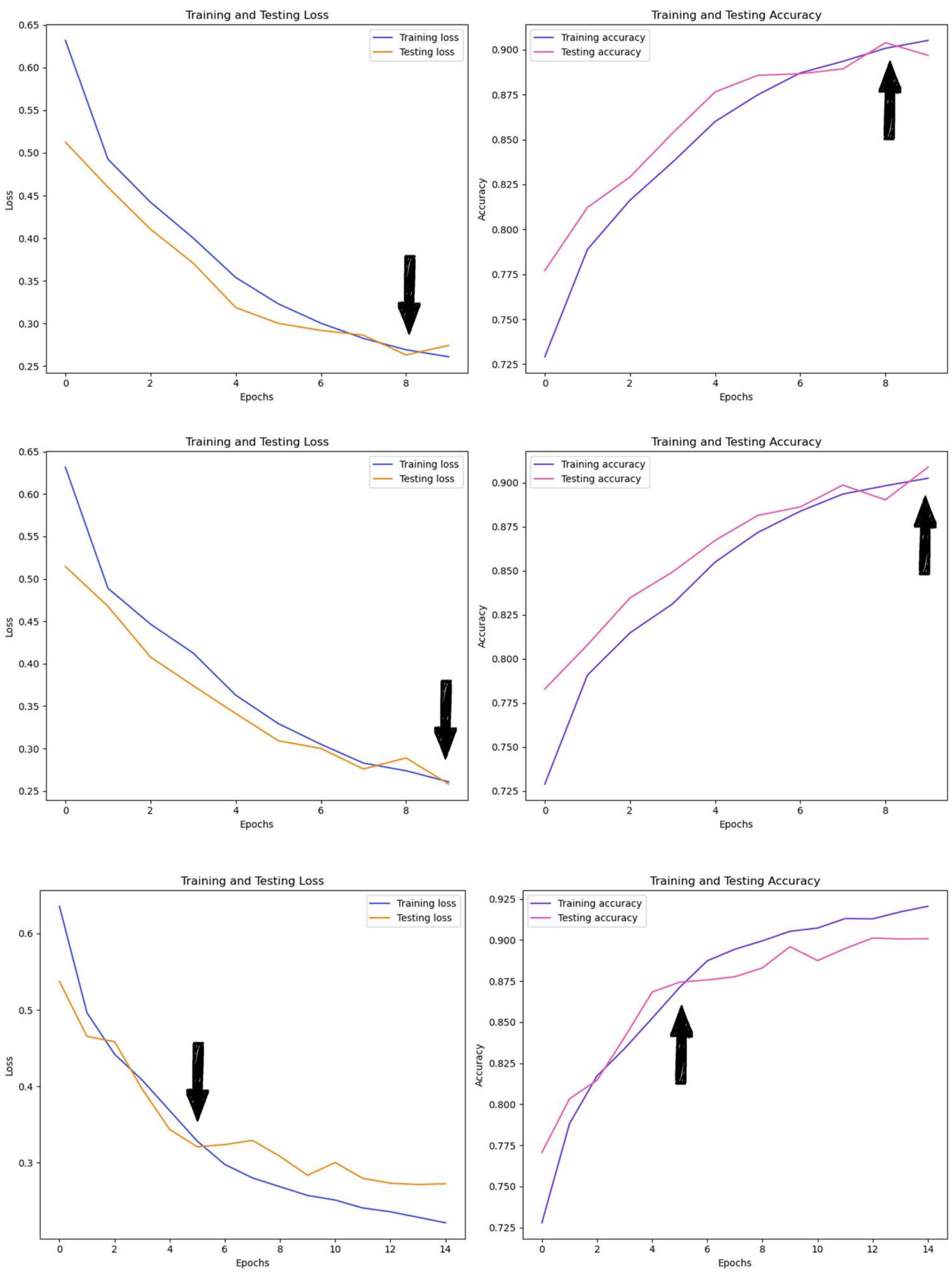


Supplementary figure 5: Every 4 mutation sequences voted by the ensemble models as "++"

A

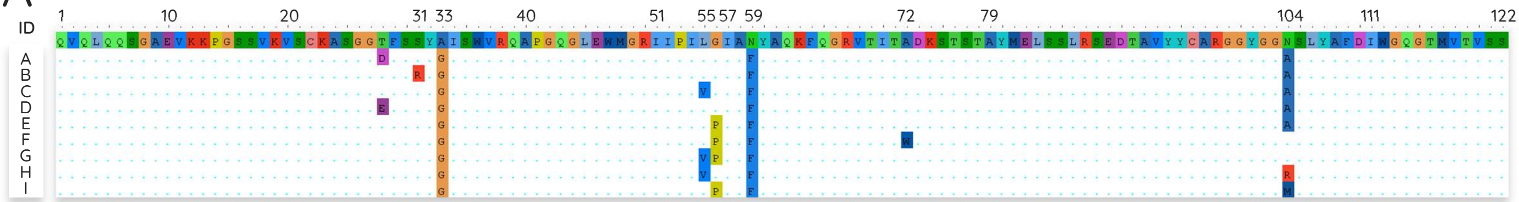


B

| ID | Ensemble Model prediction | Model A prediction | Confidence on class 0 | Confidence on class 1 | Confidence on class 2 | Model B prediction | Confidence on class 0 | Confidence on class 1 | Confidence on class 2 | Model C prediction | Confidence on class 0 | Confidence on class 1 | Confidence on class 2 | Nb mut |
|---|---|---|---|---|---|---|---|---|---|---|---|---|---|---|
| A | 2 | 1 | 3% | 59% | 37% | 2 | 1% | 34% | 65% | 2 | 3% | 32% | 66% | 4 |
| B | 2 | 1 | 2% | 76% | 22% | 2 | 1% | 42% | 57% | 2 | 3% | 40% | 58% | 4 |
| C | 2 | 2 | 1% | 16% | 83% | 2 | 0% | 6% | 93% | 2 | 1% | 10% | 88% | 4 |
| D | 2 | 1 | 4% | 66% | 30% | 2 | 2% | 35% | 62% | 2 | 4% | 34% | 62% | 4 |
| E | 2 | 2 | 3% | 36% | 60% | 2 | 1% | 19% | 80% | 2 | 1% | 9% | 90% | 4 |
| F | 2 | 2 | 1% | 32% | 67% | 1 | 5% | 53% | 42% | 2 | 5% | 38% | 57% | 4 |
| G | 2 | 2 | 2% | 34% | 64% | 1 | 7% | 50% | 43% | 2 | 8% | 27% | 65% | 4 |
| H | 2 | 2 | 2% | 41% | 57% | 2 | 1% | 17% | 82% | 2 | 2% | 23% | 75% | 4 |
| I | 2 | 1 | 4% | 57% | 39% | 2 | 2% | 33% | 65% | 2 | 2% | 21% | 77% | 4 |

### Supplementary figure 6: Analysis of 3 mutations sequences

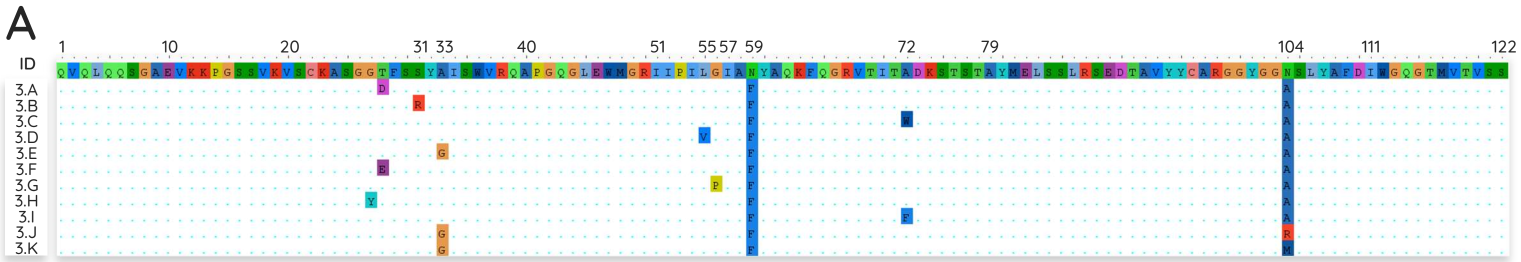


B

| ID | nb_mut | Ensemble Model prediction | Model A prediction | Confidence on class 0 | Confidence on class 1 | Confidence on class 2 | Model B prediction | Confidence on class 0 | Confidence on class 1 | Confidence on class 2 | Model C prediction | Confidence on class 0 | Confidence on class 1 | Confidence on class 2 |
|---|---|---|---|---|---|---|---|---|---|---|---|---|---|---|
| 3.A | 3 | 1 | 1 | 4% | 95% | 1% | 1 | 1% | 98% | 2% | 1 | 5% | 91% | 4% |
| 3.B | 3 | 1 | 1 | 3% | 97% | 0% | 1 | 1% | 99% | 1% | 1 | 4% | 95% | 1% |
| 3.C | 3 | 1 | 1 | 1% | 99% | 0% | 1 | 0% | 99% | 0% | 1 | 2% | 97% | 1% |
| 3.D | 3 | 1 | 1 | 3% | 94% | 4% | 1 | 1% | 89% | 10% | 1 | 6% | 85% | 10% |
| 3.E | 3 | 1 | 1 | 3% | 89% | 9% | 1 | 1% | 72% | 27% | 1 | 4% | 68% | 28% |
| 3.F | 3 | 1 | 1 | 4% | 95% | 1% | 1 | 1% | 97% | 1% | 1 | 7% | 90% | 3% |
| 3.G | 3 | 1 | 1 | 7% | 91% | 3% | 1 | 2% | 93% | 5% | 1 | 7% | 81% | 13% |
| 3.H | 3 | 1 | 1 | 5% | 94% | 0% | 1 | 1% | 99% | 0% | 1 | 6% | 93% | 1% |
| 3.I | 3 | 1 | 1 | 1% | 99% | 0% | 1 | 0% | 100% | 0% | 1 | 2% | 97% | 1% |
| 3.J | 3 | 1 | 1 | 2% | 96% | 2% | 1 | 1% | 89% | 10% | 1 | 4% | 83% | 13% |
| 3.K | 3 | 1 | 1 | 2% | 94% | 4% | 1 | 1% | 85% | 14% | 1 | 4% | 83% | 13% |

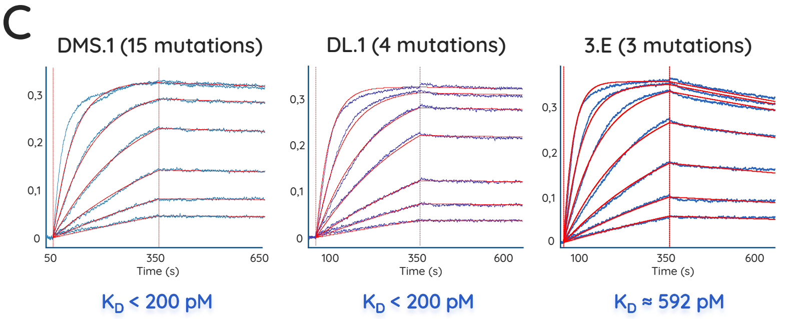


### Supplementary figure 7: Every experimental KD value compared to model predictions of antibody behavior

| clone | mutations | predicted category (vote) | model A prediction | model A Confidence | model B prediction | model B Confidence | model C prediction | model C Confidence | *in vitro* $K_D$ on mPD-L1 (BLI) |
|---|---|---|---|---|---|---|---|---|---|
| WT | 0 | **0** | 0 | 94% | 0 | 98% | 0 | 90% | 1,86E-07 |
| HM.1 | 1 | **0** | 0 | 93% | 0 | 98% | 0 | 89% | 1,47E-07 |
| HM.2 | 2 | **0** | 0 | 82% | 0 | 97% | 0 | 80% | 1,04E-07 |
| HM.3 | 3 | **1** | 1 | 96% | 1 | 68% | 1 | 69% | 2,97E-09 |
| HM.5 | 5 | **1** | 2 | 51% | 1 | 76% | 1 | 71% | 6,13E-10 |
| HM.6 | 6 | **1** | 1 | 63% | 1 | 87% | 1 | 66% | 9,47E-10 |
| HM.7 | 7 | **2** | 2 | 99% | 2 | 99% | 2 | 98% | 3,52E-10 |
| SM.1 | 15 | **2** | 2 | 100% | 2 | 100% | 2 | 100% | 2,33E-10 |
| SM.2 | 13 | **2** | 2 | 99% | 2 | 98% | 2 | 97% | 3,99E-11 |
| SM.3 | 13 | **2** | 2 | 100% | 2 | 100% | 2 | 100% | 1,14E-10 |
| SM.4 | 14 | **2** | 2 | 100% | 2 | 100% | 2 | 100% | 1,84E-10 |
| SM.5 | 14 | **2** | 2 | 100% | 2 | 100% | 2 | 100% | 2,89E-11 |
| SM.6 | 14 | **2** | 2 | 100% | 2 | 99% | 2 | 99% | 1,18E-10 |
| SM.7 | 14 | **2** | 2 | 89% | 2 | 90% | 2 | 80% | 4,20E-10 |
| SM.8 | 12 | **2** | 2 | 56% | 2 | 69% | 1 | 84% | 3,69E-10 |
| SM.9 | 12 | **2** | 2 | 93% | 2 | 99% | 2 | 79% | 2,20E-10 |
| SM.10 | 13 | **2** | 2 | 100% | 2 | 100% | 2 | 100% | 2,65E-10 |
| 3.E | 3 | **1** | 1 | 89% | 1 | 72% | 1 | 68% | 5,92E-10 |
| DL.1 | 4 | **2** | 2 | 83% | 2 | 93% | 2 | 88% | 2,29E-10 |
| DL.2 | 4 | **2** | 2 | 60% | 2 | 80% | 2 | 90% | 1,82E-10 |
| DL.3 | 5 | **2** | 2 | 96% | 2 | 98% | 2 | 97% | 1,80E-10 |
| DL.4 | 5 | **2** | 2 | 96% | 2 | 99% | 2 | 98% | 2,89E-10 |
| DL.5 | 5 | **2** | 2 | 95% | 2 | 98% | 2 | 96% | 2,35E-10 |
| DL.6 | 5 | **2** | 2 | 93% | 2 | 98% | 2 | 96% | 2,00E-10 |
| DL.7 | 5 | **2** | 2 | 94% | 2 | 95% | 2 | 99% | 2,05E-10 |
| DL.8 | 5 | **2** | 2 | 91% | 2 | 96% | 2 | 93% | 1,59E-10 |

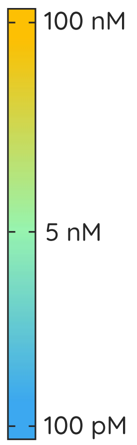

### Supplementary figure 8: DMS on PD-L1 for epitope mapping heatmaps

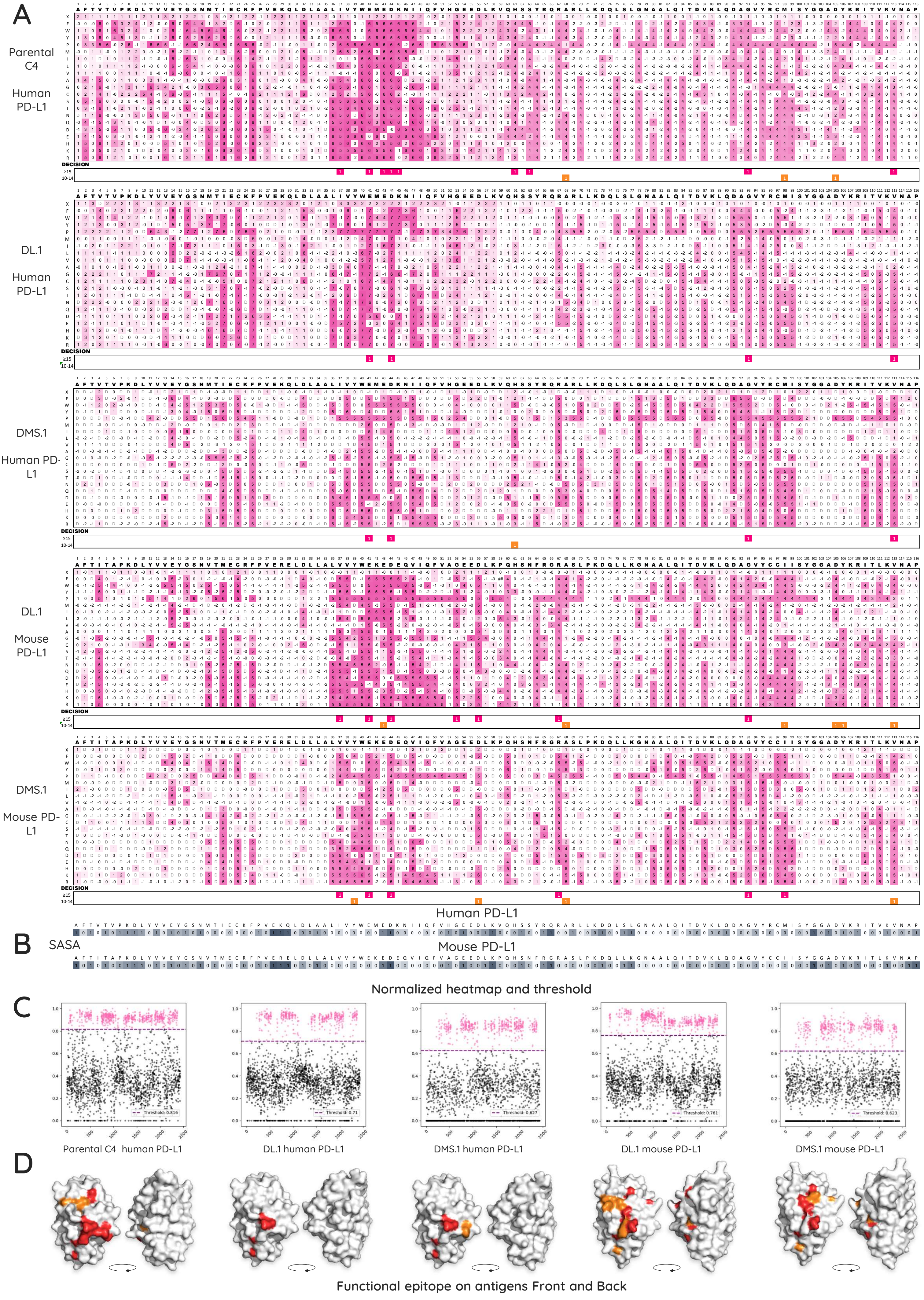

Supplementary Figure 9: Structural analysis of species-specific recognition determinants in the C4 / PD-L1 interface.

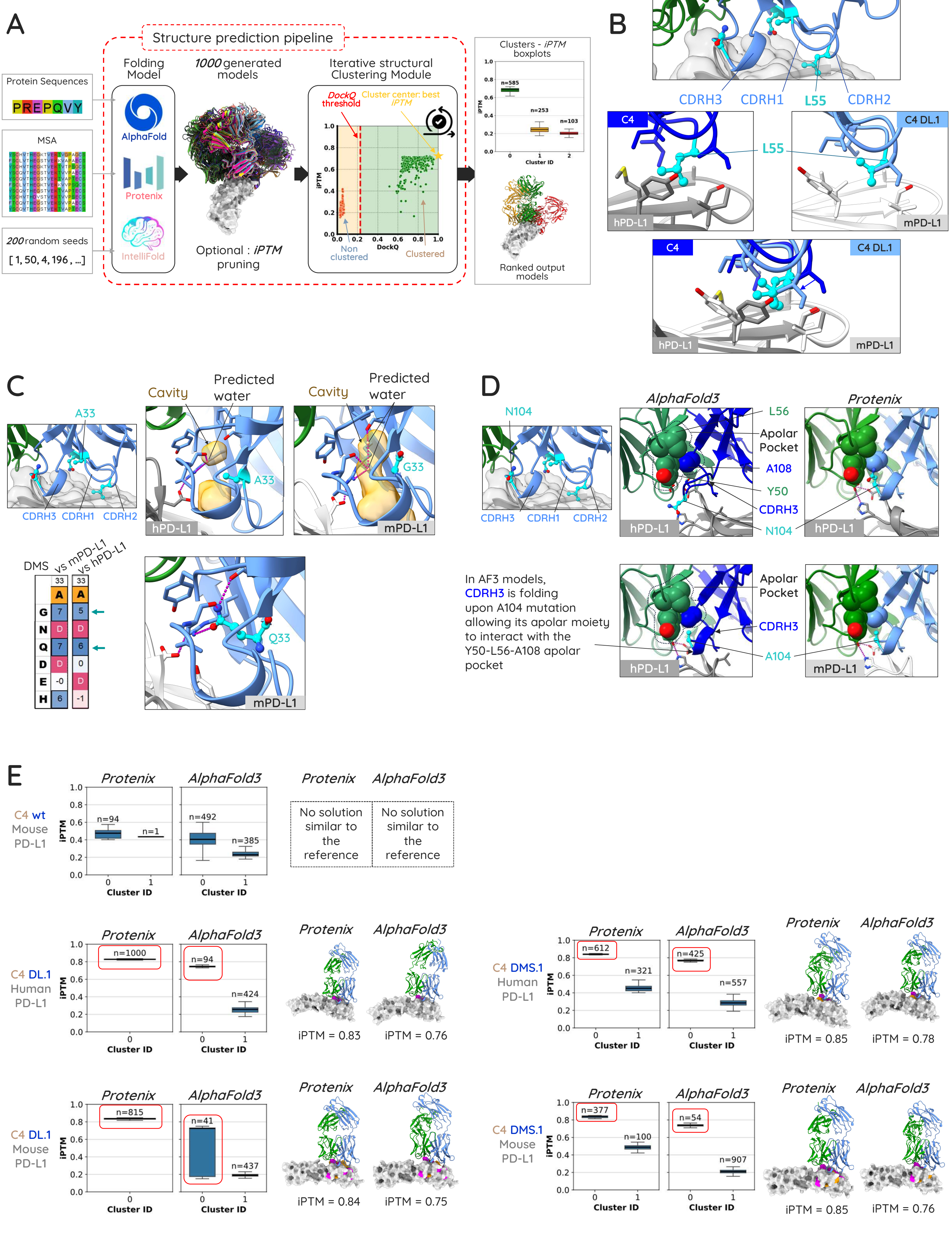
